# Brd4-bound enhancers drive cell intrinsic sex differences in glioblastoma

**DOI:** 10.1101/199059

**Authors:** Najla Kfoury, Zongtai Qi, Briana C Prager, Michael N Wilkinson, Lauren Broestl, Kristopher C Berrett, Arnav Moudgil, Sumithra Sankararaman, Xuhua Chen, Jay Gertz, Jeremy Naftali Rich, Robi D Mitra, Joshua B Rubin

## Abstract

Sex can be an important determinant of cancer phenotype, and exploring sex-biased tumor biology holds promise for identifying novel therapeutic targets and new approaches to cancer treatment. In an established isogenic murine model of glioblastoma, we discovered correlated transcriptome-wide sex differences in gene expression, H3K27ac marks, large Brd4-bound enhancer usage, and Brd4 localization to Myc and p53 genomic binding sites. These sex-biased gene expression patterns were also evident in human glioblastoma stem cells (GSCs). These observations led us to hypothesize that Brd4-bound enhancers might underlie sex differences in stem cell function and tumorigenicity in GBM. We found that male and female GBM cells exhibited opposing responses to pharmacological or genetic inhibition of Brd4. Brd4 knockdown or pharmacologic inhibition decreased male GBM cell clonogenicity and *in vivo* tumorigenesis, while increasing both in female GBM cells. These results were validated in male and female patient-derived GBM cell lines. Furthermore, analysis of the Cancer Therapeutic Response Portal of human GBM samples segregated by sex revealed that male GBM cells are significantly more sensitive to BET inhibitors than are female cells. Thus, for the first time, Brd4 activity is revealed to drive a sex differences in stem cell and tumorigenic phenotype, resulting in diametrically opposite responses to BET inhibition in male and female GBM cells. This has important implications for the clinical evaluation and use of BET inhibitors.

**Significance:** Consistent sex differences in incidence and outcome have been reported in numerous cancers including brain tumors. GBM, the most common and aggressive primary brain tumor, occurs with higher incidence and shorter survival in males compared to females. Brd4 is essential for regulating transcriptome-wide gene expression and specifying cell identity, including that of GBM. We report that sex-biased Brd4 activity drive sex differences in GBM and render male and female tumor cells differentially sensitive to BET inhibitors. The observed sex differences in BETi treatment strongly indicate that sex differences in disease biology translate into sex differences in therapeutic responses. This has critical implications for clinical use of BET inhibitors further affirming the importance of inclusion of sex as a biological variable.

---

Until recently, most basic and clinical research focused on investigating factors that influence disease susceptibility and progression without regard to biologic sex. However, mounting evidence has revealed sex differences in the incidence, age of onset, and outcome of numerous human diseases, including cardiovascular diseases, metabolic diseases, asthma, autoimmune diseases, birth defects, neurological diseases, psychiatric disorders, and cancers (1–4). The preponderance of sex differences in disease incidence and outcome led to the implementation of new guidelines by the NIH regarding inclusion of sex as a biological variable in all research.

Glioblastoma (GBM), the most common, aggressive, and incurable form of primary brain cancer (5, 6), is more prevalent in males, regardless of race or region of the world (male to female incidence of 1.6:1) (7–10). This sex difference extends across species; a male prevalence also occurs in spontaneous GBM in large dogs, suggesting a fundamental effect of sex on GBM risk (11). In addition to the sex difference in GBM incidence, Ostrom et al. documented a survival advantage for female GBM patients (12). The reasons for these sex differences in GBM incidence and outcome are largely unknown. We hypothesized that studying sex differences in GBM biology will inform the mechanisms underlying cancer risk and progression, with the ultimate goal of incorporating sex-informed approaches to treatment to improve survival of all patients.

While disease-related sex differences are often mediated through acute sex hormone actions, sex differences in the rates of multiple brain tumors are evident at all ages, as well as in neutered and non-neutered dogs (8), suggesting that factors other than circulating sex hormones underlie this skewing (13). Such factors may include the organizational or epigenetic effects of transient *in utero* sex hormones, the effects of X chromosome alleles that escape inactivation or the extra-gonadal expression of non-pseudoautosomal Y chromosome encoded genes (14–17).

We previously discovered that a genetically engineered model of GBM, involving combined loss of neurofibromin (NF1) and p53 function in murine neocortical post-natal day 1 (p1) astrocytes (male and female GBM astrocytes), exhibits sex differences *in vivo* (18, 19), including proliferation, clonogenic stemlike frequency and *in vivo* tumorigenesis, cell cycle regulation (18, 19), gene expression (18), and chemotherapy response (18) that mimic those observed in GBM patients (18, 19). Sex differences in the tumorigenic phenotype were validated in a second CRISPR-IUE (*in utero* electroporation) murine model of GBM. In this model, guide sequence inserts targeting Nf1 and p53 were injected into the lateral ventricles of embryonic pups of male and female mice, and progenitor cells were targeted via bioelectroporation. Both male and female mice developed tumors; however, male mice exhibited an accelerated tumorigenic phenotype with a shorter median survival (18). Together, these human and mouse data suggest that male and female GBM cells are differentially vulnerable to oncogenic events and to the cytotoxic effects of chemotherapy.

What are the factors that control these differences between male and female cells? In a recent multi-institutional study, we found that longer survival for male and female GBM patients after standard surgery, radiation and chemotherapy is dependent upon different transcriptional programs (20). We discerned multiple sex-specific transcriptional subtypes of GBM that correlated with survival in a sex-specific manner. We hypothesized that these sex-specific transcriptional states arise through sex-specific epigenetics, analogous to mechanisms that drive normal sexual differentiation (21, 22).

Bromodomain and extra-terminal (BET) family proteins are epigenetic readers of histone lysine acetylation and function as co-activators or corepressors of gene expression by recruiting specific transcriptional complexes to target genes (23). The BET family member Brd4 is essential for regulating transcriptome-wide gene expression and specifying cell identity, including that of GBM (24–26). Additionally, Brd4 is an emerging drug target for epigenetic interventions (23–34). Here, we show that sex differences in tumor phenotype are dependent on differential Brd4-bound enhancer regulation of stem cell-like phenotypes in male and female murine and human GBM cells. *In vitro* and *in vivo* studies of genetic or pharmacological Brd4 inhibition, as well as data mining of the Cancer Therapeutics Response Portal, indicate that male and female murine and human GBM cells are differentially sensitive to inhibition of Brd4, resulting in decreased *in vitro* clonogenicity and *in vivo* growth of male tumors, and opposite effects on female cells and tumors. Understanding the extent to which sex differences in GBM are mediated by epigenetic mechanisms will be imperative to understanding the biology of sex differences in GBM, stratifying patients for treatment with epigenetic agents, and anticipating potential sex-biased toxicities following systemic disruption of epigenetics.

## Results

### Male and female GBM cells utilize different sets of Brd4-bound enhancers

Previously, we reported correlations between sex differences in tumorigenicity and gene expression in our murine model of GBM (male and female GBM astrocytes lacking function of both NF1 and p53, see SI Materials and Methods for full details) (18, 19). Fifty percent of the sex-biased differences in gene expression observed in this model were concordantly sex-biased in human GBM expression data (18). This concordance demonstrated that our murine model recapitulates important transcriptional pathways that govern clinically relevant sex-differences in human GBM. In addition, we found correlations between sex differences in survival and gene expression in patients with GBM (20). We therefore determined whether these distinct GBM transcriptional states were the result of sexspecific enhancer activity, and whether such states could be perturbed with small molecules that target epigenetic regulators that bind at these enhancers. We prioritized Brd4-bound enhancers for analysis because these enhancers play key roles in establishing cell identity (25, 30–34), and may regulate the cell intrinsic sex differences observed in GBM. Brd4 is an epigenetic reader that binds acetylated histones H3 and H4 throughout the entire cell cycle and is deregulated in numerous cancers (35). Brd4 promotes epithelial-to-mesenchymal transition, stem cell-like conversion, and pluripotency (26, 28, 36). Brd4 pharmacological inhibition has shown therapeutic activity in a number of different cancer models (37–41).

To evaluate a potential role for Brd4 in mediating sex differences in GBM, we mapped Brd4 genomic localization in male and female GBM astrocytes (highly active Brd4-bound enhancers and typical Brd4-bound enhancers) using transposon Calling Cards (42, 43) to identify enhancers differentially bound by Brd4. To do so, we fused the piggyBac (PB) transposase to the C terminus of the Brd4 protein, endowing the transposase with the ability to direct the insertion of the PB transposon into the genome close to Brd4 binding sites. Five biological replicates were carried out, and the correlation between replicates was r > 0.9 for all pairwise comparisons (*SI Appendix,* Fig. 1*A*). Using this protocol, we mapped ~1.25 million unique insertions directed by the Brd4-PBase fusion for each male and female sample. As Brd4 is an epigenetic reader of the acetylated lysine 27 residue of histone H3 (H3K27ac), we confirmed that these newly identified Brd4-bound enhancers were enriched for H3K27ac. We performed H3K27ac Chromatin Immunoprecipitation Sequencing (ChIP-seq) in male and female GBM cells to identify the genomic regions enriched for this well-known marker of active enhancers (44). Three biological replicates were analyzed and the correlation between replicates was r > 0.9 for all pairwise comparisons (*SI Appendix,* Fig. 1*B*). As expected, data collected from calling cards and H3K27ac ChIP-seq revealed a high concordance between Brd4-bound enhancers and H3K27ac enrichment. A representative example of concordant Brd4 binding and H3K27ac in male and female GBM astrocytes at high and low-enriched regions is depicted in Fig. 1*A*. We next analyzed the distances from Brd4 binding sites to the nearest H3K27ac-enriched regions. Brd4 binding sites were significantly enriched at H3K27ac candidate enhancer regions (Fig. 1*B*) and display concordant gene expression levels (Fig. 1*C*) indicative of active Brd4-bound enhancers *(p* < 0.01). More specifically, 82% and 81% of Brd4 bound enhancers were localized within 200bp of H3K27ac peaks in male and female GBM cells, respectively.

**Fig. 1.**
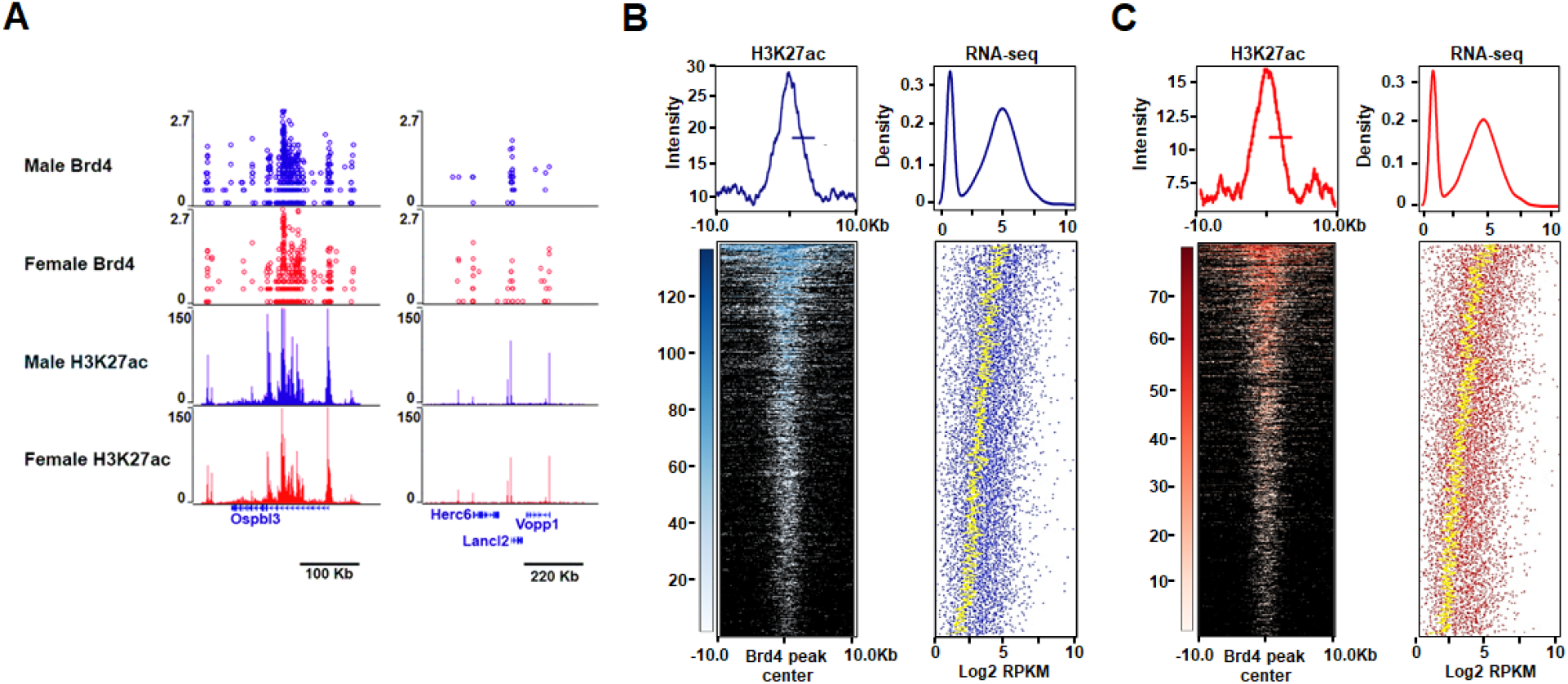
Concordance between Brd4-bound enhancers, H3K27ac enrichment, and gene expression in mouse GBM astrocytes. (*A*) A representative example of concordant Brd4 and H3K27ac binding at high and low-enriched regions in male and female GBM astrocytes. (*B*, *C)* Heatmaps and intensity plots of H3K27ac signal at sex-biased Brd4-enriched peaks (+/− 10 Kb from the Brd4 peak center) in male (B) and female (*C*) GBM cells. For each sex-biased Brd4-enriched peak, the expression values of the nearest gene are presented in a dot plot in which the yellow-dotted line depicts the average gene expression levels of 50 genes (Density refers to the fraction of genes). These analyses revealed that Brd4 binding sites are significantly enriched at H3K27ac candidate enhancer regions and display concordant gene expression levels indicative of active Brd4-bound enhancers (p < 0.01).

Having established that our genomic data was reproducible and that Brd4 binding sites occur at genomic regions enriched for H3K27ac, we next sought to identify the genomic loci that were differentially bound by Brd4 in male and female cells. We identified 2679 enhancers (20% of all Brd4-bound enhancers in male GBM cells) that bound more Brd4 protein in males and 2778 enhancers (21.2% of all Brd4-bound enhancers in female GBM cells) that bound more Brd4 protein in females (Fig. 2*A*). The Brd4 signal intensity for these male-biased and female-biased enhancers in male and female GBM astrocytes is depicted in Fig. 2*B*. To validate these putative male- and female-biased enhancers using an orthogonal method, we analyzed their H3K27ac status and found that the loci that bound Brd4 in a sex-biased manner displayed sex-biased enrichment of H3K27ac (Fig. *2C*). Chromosome location analysis for these sex-biased Brd4-bound enhancers revealed that only 0.13% and 3.11% of male-biased Brd4-bound enhancers were located on the Y and X chromosomes, respectively, and that only 4.29% of female-biased Brd4-bound enhancers were located on the X chromosome, indicating that the observed differences in Brd4-bound enhancers are not simply due to differential Brd4 enhancer enrichment on sex chromosomes. The majority of enhancers exhibited a sex-bias in Brd4 occupancy, while a subset appeared to be sex-specific. We will collectively refer to these enhancers as sex-biased. Representative examples of sex-specific, Nkx2.1, and sex-biased, Zic1/4, enhancers, are depicted in Fig. 2 *D* and *E*. Nkx2.1 functions both as a “lineage survival” oncogene and tumor suppressor in lung adenocarcinomas, (45, 46) whereas Zic1 acts as a tumor suppressor in breast and thyroid cancers (47, 48), supporting the potential context-dependent dual role of Brd4 in oncogenesis. This is the first demonstration of differential Brd4-bound enhancer usage by male and female cells of any kind, suggesting that these enhancers may function in sexual differentiation in a manner similar to their role in determining cell identity and fate (25, 30–32, 49). A violin plot illustrating the full distribution of the enrichment signal for male and female-biased loci and shared peaks is depicted in Fig. *2F*.

**Fig. 2.**
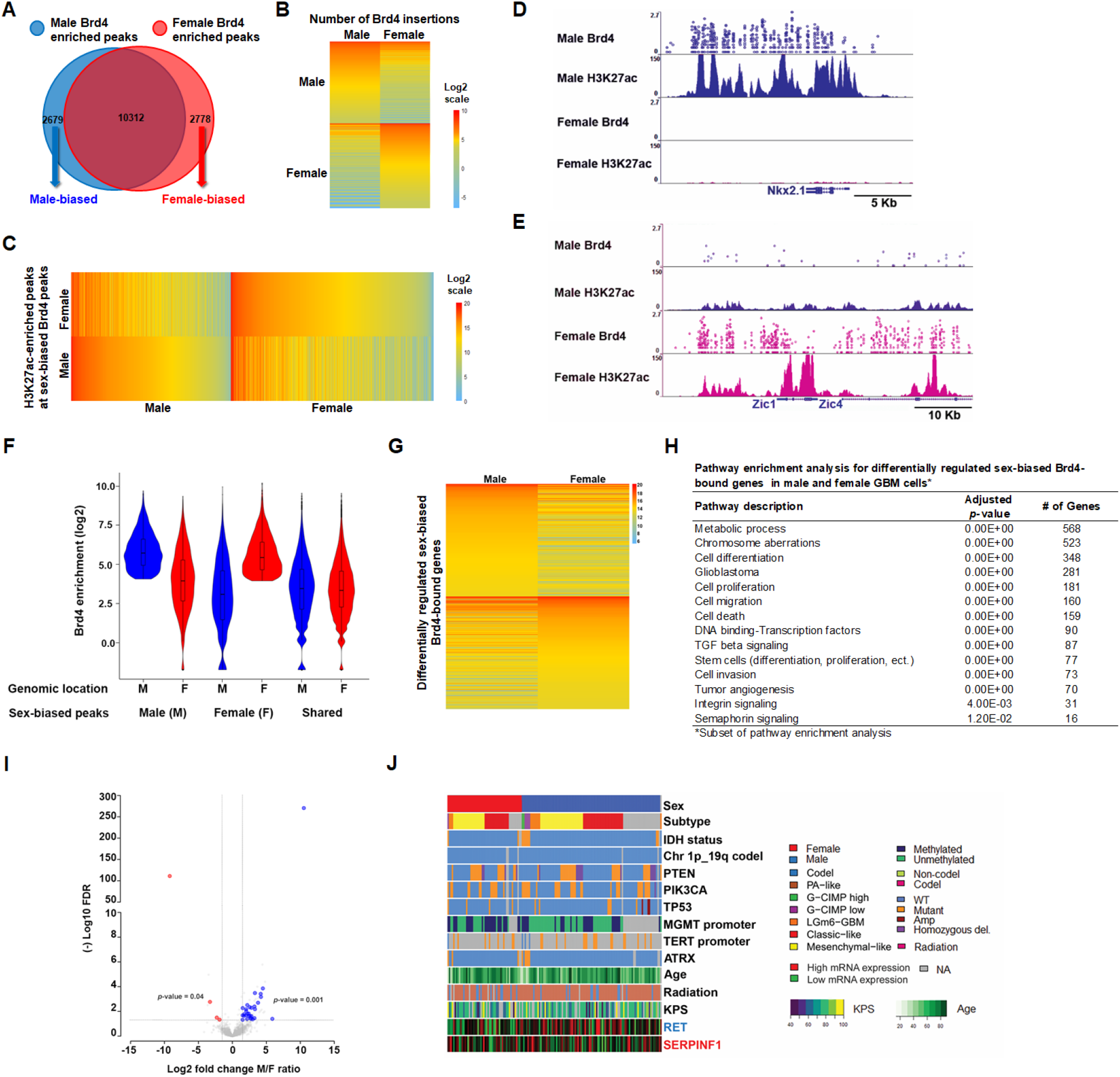
Male and female GBM cells have sex-biased Brd4-bound enhancers. (*A*) Venn-diagram showing the number of unique male (2679), unique female (2778), and overlapping (10,312) Brd4-enriched peaks identified in GBM astrocytes. *(B)* A heatmap depiction of the number of Brd4 insertions at male and female-biased Brd4-enriched peaks in GBM astrocytes (normalized data). (*C*) A heatmap analysis of H3K27ac signal intensity (read depth) at male and female-biased Brd4-enriched peaks in GBM astrocytes. *(D)* Nkx2.1 and *(E)* Zic1/4 illustrate male-specific and female-biased genes, respectively, associated with differential Brd4 binding affinity and H3K27ac enrichment. The x-axis (blue arrows) of all tracks corresponds to genomic location of the gene. The y-axis of Calling Card tracks represents the log10 scale of sequencing reads for each insertion as indicated by circles. The y-axis of ChIP-seq tracks represents the number of uniquely mapped reads. *(F)* Violin plot illustrating the full distribution range of the enrichment signal for male and female biased-loci and shared peaks at different genomic locations. (*G*) RNA abundance (RNA sequencing) in male and female GBM cells for differentially regulated sex-biased Brd4-bound genes (n=3). *(H)* Pathway enrichment analysis for differentially regulated sex-biased Brd4-bound genes in male and female GBM astrocytes. *(I)* Concordance in sex-specific gene expression patterns between murine GBM lines and human GSCs. Volcano plot depicting all mouse sex-biased significantly expressed genes with color-coded concordant human sex-biased genes (blue circles_male-biased genes; red circles_female-biased genes; log2FC 1.5 and FDR 0.05) genes. The x-axis represents mouse log2 fold change of male/female ratio for each significantly expressed sex-biased gene with its corresponding (-) log10 FDR value on the y-axis. *(J)* Gene expression pattern, whole-exome sequencing and clinical phenotype data from the TCGA GBM datasets for 2 representative examples of concordant human and mouse male (blue) and female (red) sex-biased genes.

We next sought to identify the genes regulated by these sex-biased Brd4-bound enhancers. It is challenging to reliably link enhancers to the genes they regulate (50), so we used conservative criteria: we linked enhancers to the nearest gene within a 5kb genomic distance. This analysis revealed 1740 male-biased genes and 1604 female-biased genes. Pathway enrichment analysis using Genomatix Pathway System (GePS) of male-biased enhancer-gene pairs revealed functional enrichment for glioblastoma, cell proliferation, cell cycle, regulation of transcription, tumor angiogenesis and cancer stem cells. Similar analysis on female-biased enhancer-gene pairs showed an enrichment in pathways involved in cell differentiation, glioblastoma, regulation of transcription, cell migration and neural stem cells (*SI Appendix,* Table 1).

Based on our observation of sex-biased Brd4-bound enhancer usage between male and female GBM cells, we profiled male and female GBM cells with RNA sequencing (RNA-seq). For each condition, three biological replicates were performed; the data were highly reproducible (Pearson r ≥ 0.96 for all pairwise comparisons) (*SI Appendix,* Fig. 1*C*) and indicated differential expression of 3490 transcripts (FDR < 0.05). We next integrated the male and female-biased Brd4 regulated genes with the differentially expressed genes between male and female GBM astrocytes and narrowed the list of differentially regulated sex-biased Brd4-bound genes to 1296, of which 52.4% were male-biased and 47.6% were female-biased Brd4-regulated genes. A heatmap depiction of the top 400 differentially regulated sex-biased Brd4-bound genes is presented in Fig. *2G*. Pathway enrichment analysis for the 1296 differentially regulated sex-biased Brd4-bound genes was performed using a combination of KEGG pathway and Genomatix Pathway System (GePS). Classification of these genes according to function revealed a significant number of relevant and important pathways, including cell differentiation, glioblastoma, cell proliferation, cell migration and invasion, tumor angiogenesis, stem cell functions and DNA binding-transcription factors (Fig. *2H* and *SI Appendix*, Table 2). Thus, these sex-specific transcriptomic differences in core cancer pathways likely arise from male and female-biased Brd4-bound enhancer activity.

We next sought to determine whether these sexspecific transcriptome differences are also present in human GBM samples. To do so, we re-analyzed published RNA-seq data of 43 human glioma stemlike cells (GSCs) and segregated them by sex (51). We found 246 differentially regulated genes between male and female GSCs, of which 223 were upregulated in male and 23 were upregulated in females (log2FC 1.5 and FDR 0.05). We then compared the gene lists of sex-biased differentially regulated genes in human GSCs with those of the murine GBM astrocytes at the same cut-off parameters (log2FC 1.5 and FDR 0.05; 981 and 771 upregulated genes in male and female GBM cells, respectively) and found an overlap in gene identity and direction of change for both males (17.5% of human male-biased differentially expressed genes, *p-*value 0.001) and females (17.4% of human female-biased differentially expressed genes, *p-*value 0.04) (*SI Appendix*, Table 3). *p-*values were calculated using the cumulative hypergeometric distribution. A volcano plot depicting all mouse sex-biased significantly expressed genes with color-coded concordant human sex-biased genes (blue circles_male-biased genes; red circles_female-biased genes; log2FC 1.5 and FDR 0.05) genes is presented in Fig. 2*I*. The expression patterns of two representative examples of concordant male-biased (RET) and female-biased genes (SERPINF1) are illustrated in Fig. *2J*, with gene expression (RNA-seq), whole exome and clinical phenotype data collected from TCGA GBM data (52). Pathway enrichment analysis for the human and mouse concordant genes identified functional enrichment for the MAPK family signaling cascade, cell adhesion, mesenchymal and pluripotent stem cells, neurofibromatosis, neuroendocrine tumors and neuron differentiation. Altogether, the mouse and human data suggest the presence of sex-specific transcriptional programs in tumorigenesis that are regulated by Brd4-bound enhancers in a sex-biased manner.

### Depletion or inhibition of the BET family protein Brd4 has opposing effects on clonogenicity in mouse and human male and female GBM cells

Our genomic analyses of mouse and human GBM cells suggested to us that Brd4-bound enhancers might regulate a large number of genes and transcriptional networks in a sex-biased fashion. To test this hypothesis, we performed genetic depletion of individual BET family members in our murine model of GBM. Under basal conditions, male and female GBM cells expressed Brd3 and Brd4 mRNA at similar levels, whereas Brd2 was expressed at higher levels in female cells *(*p=* 0.02) (Fig. 3*A*). We evaluated the potency of 5 shRNAs specific to each of the Brd2, Brd3, or Brd4 genes and selected the shRNAs that achieved the most robust knockdown of each gene for downstream functional experiments of the tumorigenic phenotype. Brd2, Brd3, and Brd4 mRNA levels were partially depleted in male and female GBM cells after infection with the corresponding lentiviral shRNAs (Fig. 3*B*). Knockdown of Brd2 did not affect clonogenic frequency in either male or female cells as measured by the extreme limiting dilution assay (ELDA) (Fig. 3*C*). Male GBM cells with Brd4 knockdown exhibited a decrease in clonogenic frequency, whereas female cells displayed an increase in clonogenic frequency. A similar effect on clonogenic frequency was observed following Brd3 knockdown in male cells, although it was not as robust as the effect of Brd4 knockdown. Strong Brd4 effects were observed despite the more modest knockdown of Brd4 compared to Brd3.

**Fig. 3.**
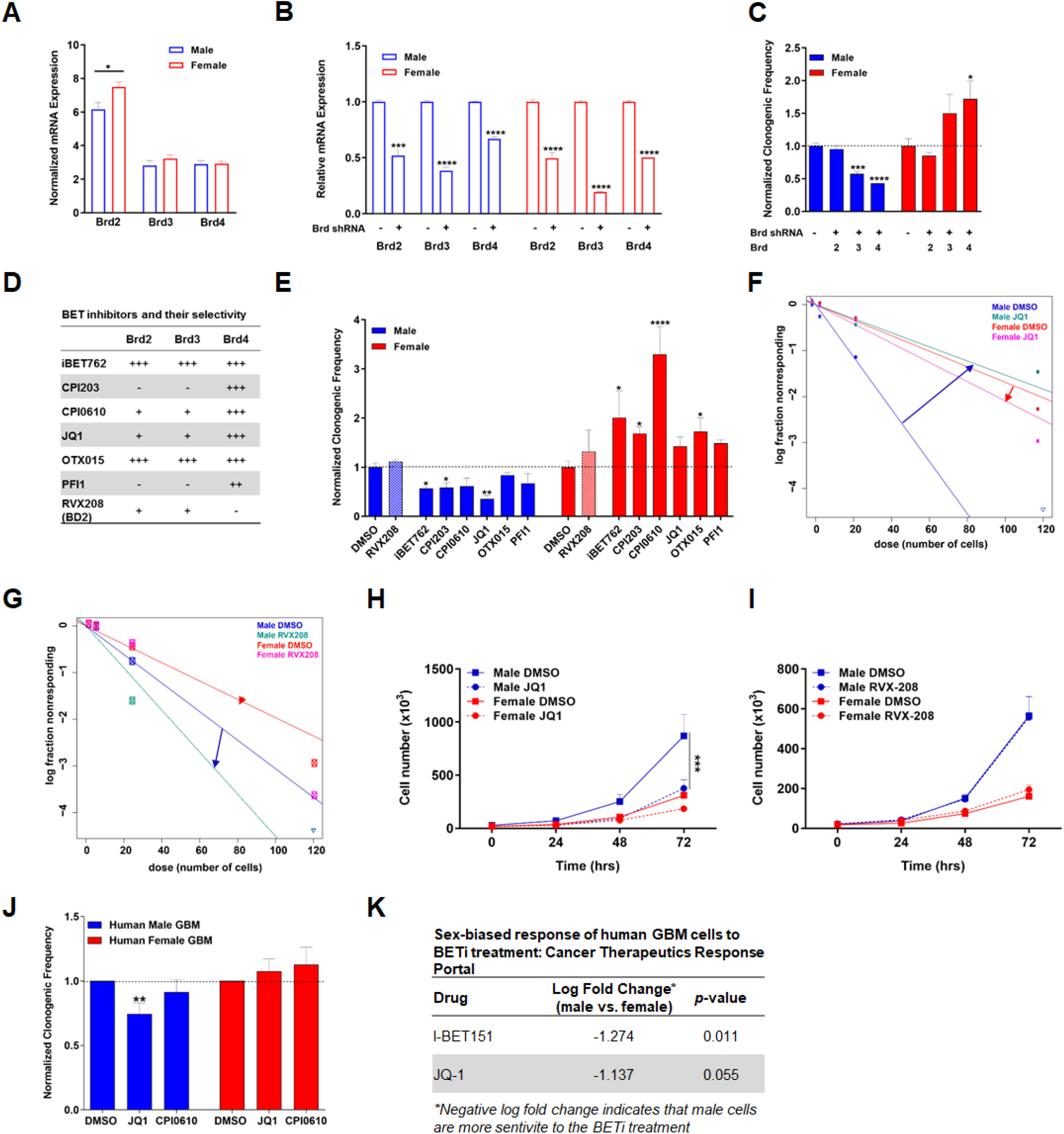
Brd4 inhibition has opposing effects on clonogenicity in mouse and human male and female GBM cells. (*A*) Normalized Brd2, Brd3 and Brd4 mRNA expression in male and female GBM cells under basal conditions. Brd3 and Brd4 are expressed at equivalent levels between male and female GBM cells, while Brd2 is expressed at higher levels in female cells. (*B*) Brd2, Brd3, and Brd4 mRNA expression following lentiviral shRNA infection. mRNA levels in knockdown samples were normalized to their respective control. (*C*) Normalized clonogenic cell frequency as determined by ELDA assay in control and Brd2, Brd3, and Brd4 knockdown male and female GBM cell lines. Knockdown of Brd4 and Brd3 suppresses clonogenic frequency in male GBM cells while female cells showed a significant increase in clonogenic frequency following Brd4 knockdown only. Brd2 depletion was without effect. All treatment groups were normalized to control clonogenic frequency levels. *(D)* Tabular representation of the BET inhibitors currently in clinical trials and their selectivity to the three BET family members. *(E)* Normalized clonogenic cell frequency as determined by ELDA assay in male and female GBM cells treated with DMSO (control) or the indicated BET inhibitor. BET inhibitors significantly reduced clonogenic cell frequency in male cells, and significantly increased clonogenic cell frequency in female cells. Shaded bars are used to indicate the effect of RVX208, a Brd2/3 inhibitor as a control for Brd4 inhibition. *(F)* Frequency of clonogenic stem-like cells as determined by ELDA assay in male and female GBM cells treated with DMSO or JQ1 (Brd2/3/4 inhibitor). Male cells exhibited greater clonogenic cell activity than female cells under control conditions. JQ1 significantly reduced clonogenic cell frequency in male cells to levels almost equivalent to female cells under control conditions, while female cells exhibited an increase in their clonogenic cell frequency. (*G*) Frequency of clonogenic stem-like cells was measured by ELDA in male and female GBM astrocytes following RVX-208 treatment, a BD2 inhibitor with selectivity to Brd2 and Brd3. No significant change in clonogenic frequency was observed in male and female GBM astrocytes following RVX208 treatment. (*H*) Proliferation assays were performed in mouse GBM cells treated with DMSO or JQ1. Male cells exhibited greater proliferation than female cells under control conditions. JQ1 significantly reduced growth in male cells to female basal growth level. (*I*) Proliferation rates of male and female GBM astrocytes were unaffected by RVX-208 treatment. (*J*) Normalized clonogenic cell frequency as determined by ELDA assay in male and female human primary GBM cells treated with DMSO (control), JQ1, or CPI0610. JQ1 treatment significantly reduced clonogenic cell frequency in human primary GBM male cells, and slightly increased clonogenic cell frequency in human primary GBM female cells. CPI0610 treatment had a similar response pattern as JQ1 in male and female human GBM primary cells. *(K)* Tabular representation of the sex-biased growth response of human GBM cells to BETi treatment. Male GBM cells are more sensitive to BETi treatment than female GBM cells (Cancer Therapeutics Response Portal). (*=*p* <0.05, **=*p* <0.01, ***=*p* <0.001, ****=*p* <0.0001 as determined by two-tailed t-test or one-way ANOVA).

Small molecule inhibitors of the BET family of proteins (BETi) are a novel class of epigenetic compounds that selectively target BET proteins and have been shown to have promise as cancer therapeutics, decreasing cell proliferation and invasion in many cancer types, including GBM (24, 27, 29). However, almost all of the American Type Culture Collection (ATCC) human cell lines used in the preclinical studies are of male origin and thus, potentially uninformative with regard to sex differences in drug effects. Therefore, we evaluated whether there was a sex difference in the treatment response to some of the BET inhibitors currently in clinical trials for cancer in our mouse GBM cells. We treated male and female GBM astrocytes with a panel of BETi (Fig. 3*D*), which are being tested in clinical trials (24 clinical trials listed in www.clinicaltrials.gov and Selleckchem.com) and then performed ELDAs to measure clonogenic cell frequency (53–55). Treatment with BETi reproducibly decreased clonogenic frequency in male GBM cells while increasing clonogenic frequency in female cells (Fig. 3 *E-G* and *SI Appendix*, Fig. 2). This striking abrogation of sex differences, driven by sex-dependent opposing response to Brd4 inhibition, is illustrated in response to the Brd4 antagonist JQ1 (Fig. *3F*). JQ1 is a thieno-triazolo-1,4-diazepine that displaces Brd4 from chromatin by competitively binding to the acetyllysine recognition pocket (37, 38). Treatment of acute myeloid leukemia cells with JQ1 causes a rapid release of Mediator 1 (MED1) from a subset of enhancer regions that were co-occupied by Brd4, leading to a decrease in the expression of neighboring genes (56). We treated male and female GBM cells with either 0.05% DMSO or 500 nM JQ1 and then performed ELDAs to measure clonogenic cell frequency. Male cells exhibited greater clonogenic cell activity than female cells under control conditions. Treatment with JQ1 reproducibly abrogated the basal differences in clonogenic frequency between male and female GBM cells by decreasing the clonogenic cell fraction in male cells and increasing the clonogenic cell fraction in female cells, rendering the male-JQ1 treated cells equivalent to the female control cells (Fig. *3F*). To confirm that these effects were mediated by Brd4-bound enhancers, we performed similar experiments using RVX-208, a BD2 inhibitor with high selectivity to Brd2/3 (57). Consistent with the Brd2 and Brd3 knockdown results, treatment with 5μM RVX-208 for 24 hours did not have an effect on the clonogenic frequency of male and female cells (Fig. 3 *E* and *G*). Next, we examined the effect of drug treatment on cell growth and proliferation. Male cells exhibited decreased proliferation and a growth phenotype equivalent to control female cells following JQ1 treatment (Fig. *3H*), whereas treatment with RVX208 did not alter proliferation of male and female cells (Fig. 3*I*). Taken together, these results demonstrate for the first time that the sex differences in the tumorigenic phenotype we observe in our murine GBM cells are mediated by differential Brd4-bound enhancers, and that the response to BET inhibition is sex-dependent.

Next, we investigated whether the sex-specific responses to BETi observed in our murine GBM model were also present in human GBM. To do so, we treated male and female human GBM primary cell lines with 0.05% DMSO, 500 nM JQ1, or 1 μM CPI0610 and then performed ELDA to measure clonogenic cell frequency. Similar to the murine data, treatment with BETi decreased clonogenic cell frequency in male patient-derived GBM cells, while slightly increasing the clonogenic cell frequency in female patient-derived GBM (Fig. *3J* and *SI Appendix,* Fig. 3). Although we observed the expected trend in response to BETi treatment in human male and female primary cell lines, our sample size was small with limited power. Additionally, human GBM samples are intrinsically heterogeneous with different genetic backgrounds, and hence the magnitude of treatment effects can be more variable compared to isogenic cell lines. To expand our sample size, we analyzed the Cancer Therapeutics Response Portal (CTRP) to examine if human male GBM cells are indeed more sensitive to BETi than female cells. First, we segregated the CTRP human GBM samples by sex. We then analyzed the cell growth response of male and female GBM cells to BETi treatment. This analysis revealed that male GBM cells are more sensitive to BETi, displaying slower growth and more cell death, consistent with what we observed using our murine GBM model (Fig. *3K*).

While the female stem cell response to BETi treatment data was variable, a clear difference in response to BET inhibition was present in the human GBM cell lines. This result was also supported by the sex difference in proliferation following BETi identified in the CTRP datasets. Thus, the sex-specific effects of Brd4 inhibition on tumorigenic phenotype and response to treatment observed in our murine GBM model extend to human GBM cells. Taken together, these results demonstrate for the first time that sex differences in GBM cellular phenotypes are mediated by differential Brd4-bound enhancers, and that reduced Brd4 function results in opposing sex-dependent effects in both mouse models and human patients. These data further affirm the sex-dependent role of Brd4 in regulating tumorigenesis in GBM and demonstrate how clinically important it will be to consider sex in the design of clinical trials and the decision making for treatment of cancer patients.

### Brd4-bound enhancers regulate sex differences in GBM

Having established that Brd4-bound enhancers mediate sex-difference in our murine model of GBM, we next sought to determine the transcriptional pathways responsible for these basal sex differences. To do so, we used our Brd4 binding data in male and female GBM astrocytes to categorize Brd4-regulated genes as male-biased (Fig. 4*A*), female-biased (Fig. *4B*), or shared (Fig. 4*C*). Although these genes were categorized using only Brd4 binding data, we found a high degree of concordance with H3K27ac intensity and gene expression within each category. When considered as a group, 95% of the male-biased Brd4-regulated genes displayed more H3K27ac signal in males and 72% were more highly expressed in males (Fig. 4*A*, *p*-value <10^-6^), while 93% of the female-biased Brd4-regulated genes displayed more H3K27ac signal in females and 67% were more highly expressed in females (Fig. 4*B*, *p*-value <10^-6^). These results demonstrate that sex-biased Brd4 binding correlated with sex-specific gene expression patterns.

**Fig. 4.**
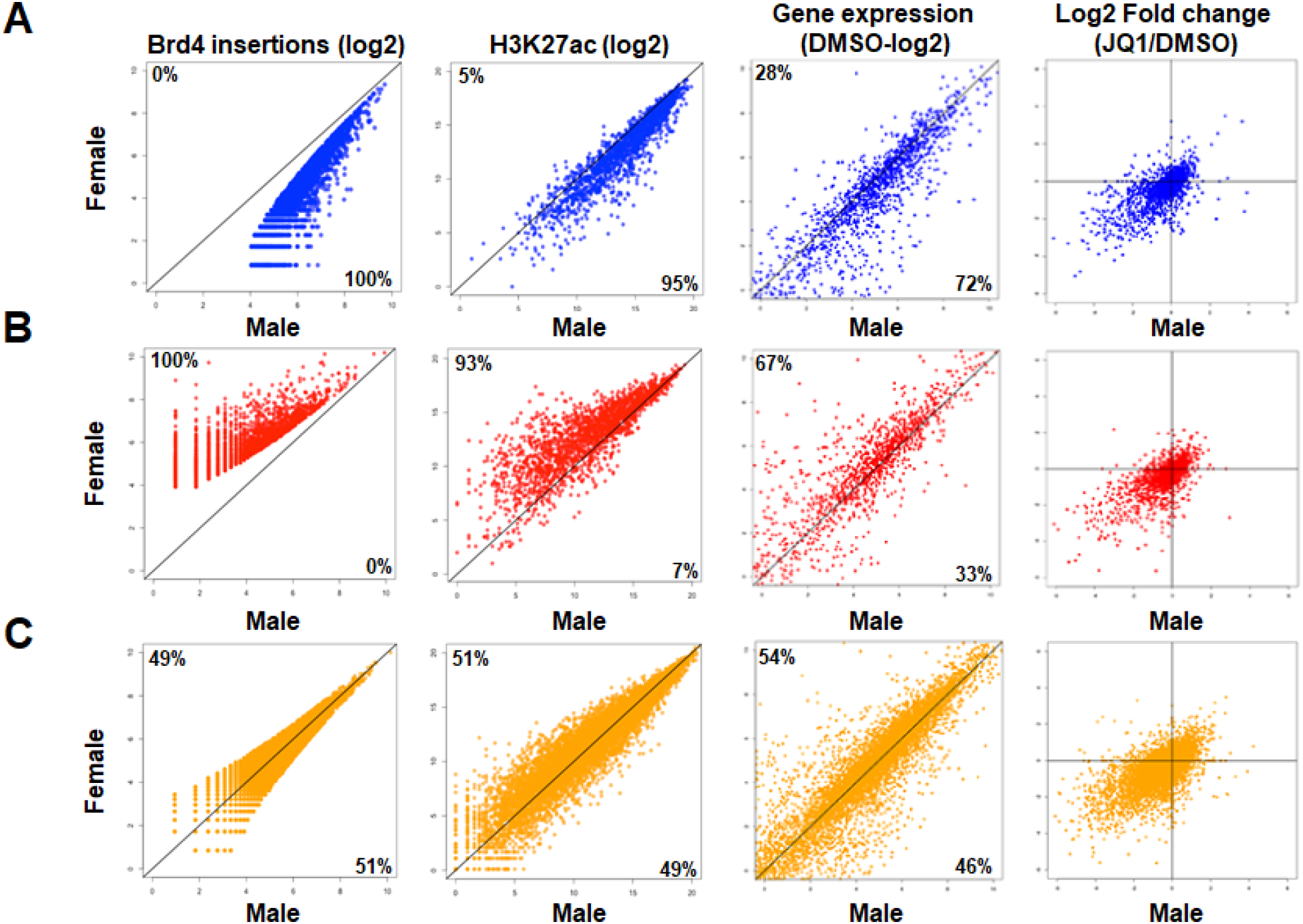
Male and female GBM cells have sex-biased Brd4-bound enhancers and concordant gene expression. Brd4 insertions, H3K27ac signal, and gene expression values (normalized Reads Per Kilobase Million (RPKM)) of the nearest gene(s) in male (x-axis) and female (y-axis) GBM astrocytes for male-biased (*A*), female-biased (*B*), and shared (*C*) genes are plotted. The percentage of genes enriched in each sex is indicated at the top left (female) and bottom right (male) of each graph. (Far right column) Log2 fold changes (Log2FC) of the gene expression value following JQ1 treatment are plotted as above. Most genes are located in the bottom, left quadrant, indicating down regulation of the gene in both sexes following JQ1.

To determine which sex-biased genes were modulated after treatment with a BET inhibitor, we performed RNA-seq on male and female GBM cells treated with JQ1, one of the BET inhibitors that had a strong sex-biased effect on clonogenicity (*SI Appendix*, Fig. 1*D*). Briefly, cells were treated with either vehicle (0.05% DMSO) or 500 nM JQ1 for 24 hours prior to RNA isolation. Gene expression analysis of JQ1- and DMSO-treated cells revealed that JQ1 treatment predominantly downregulated gene expression (Fig. 4, last column of scatterplots, and *SI Appendix*, Fig. 4*A*). In addition, genes proximal to Brd4 binding sites were significantly downregulated compared to genes that were distal to Brd4 binding sites *(p* <0.01), indicating that JQ1 had a specific and directed effect on genes whose expression was driven by Brd4-bound enhancers (*SI Appendix*, Fig. 4*B*). Pathway analysis of sex-biased Brd4-bound enhancer associated genes downregulated following JQ1 treatment in male and female GBM cells revealed functionally important pathways, such as chromosome aberrations, integrin signaling, and stem cell proliferation in males, and immune system process, cell cycle, and transforming growth factor-beta signaling in females (*SI Appendix,* Table 4).

To begin to understand the sex-specific stem cell response to BETi, we focused our attention on the functions of the sex-biased male and female Brd4-bound enhancer genes under basal conditions. Functional classification of these genes using pathway enrichment analysis revealed that male GBM cells were highly enriched for cancer, neoplastic, and pluripotent stem cell pathways, while female GBM cells were mainly enriched for neural stem cell and stem cell proliferation pathways (Fig. 5*A*). Representative examples of these pathways are depicted in Fig. 5 *B* and *C*. Male-biased genes in stem cell pathways included well-known oncogenes, such as Myc, Tet1, and Lif, while female-biased genes included tumor suppressors, such as Six2 and Six3. Based on these results and our previous observation of a sex-dependent opposing response in clonogenicity following Brd4 inhibition with JQ1, we examined the effects of JQ1 treatment on the expression of stem cell pathway genes. As BETi reduced sex differences in clonogenicity, we hypothesized that sex differences in the expression of stem cell-related genes present in male and female GBM cells under basal conditions would be eliminated by JQ1 treatment, accounting for the abrogation of sex differences in the tumorigenic phenotype. We identified genes with sex differences in expression at baseline that became equivalent following JQ1 treatment, and performed pathway analysis. As anticipated, the cancer stem cell pathway was affected by JQ1 treatment (Fig. 5*D*). Concordant with differential growth responses, JQ1 downregulated male-biased genes, while JQ1 upregulated female-biased genes. The fold change in expression following JQ1 treatment (ratio of JQ1/DMSO) is depicted above or below each gene. Overall, there was concordance with the direction of change in gene expression and the decrease and increase in stem cell function in male and female ELDA assays respectively. These results support the existence of Brd4-regulated sex-specific transcriptional program that mediate key functional properties of GBM, including clonogenicity. Identifying the specific pathways critical to sex differences in GBM will require further functional studies.

**Fig. 5.**
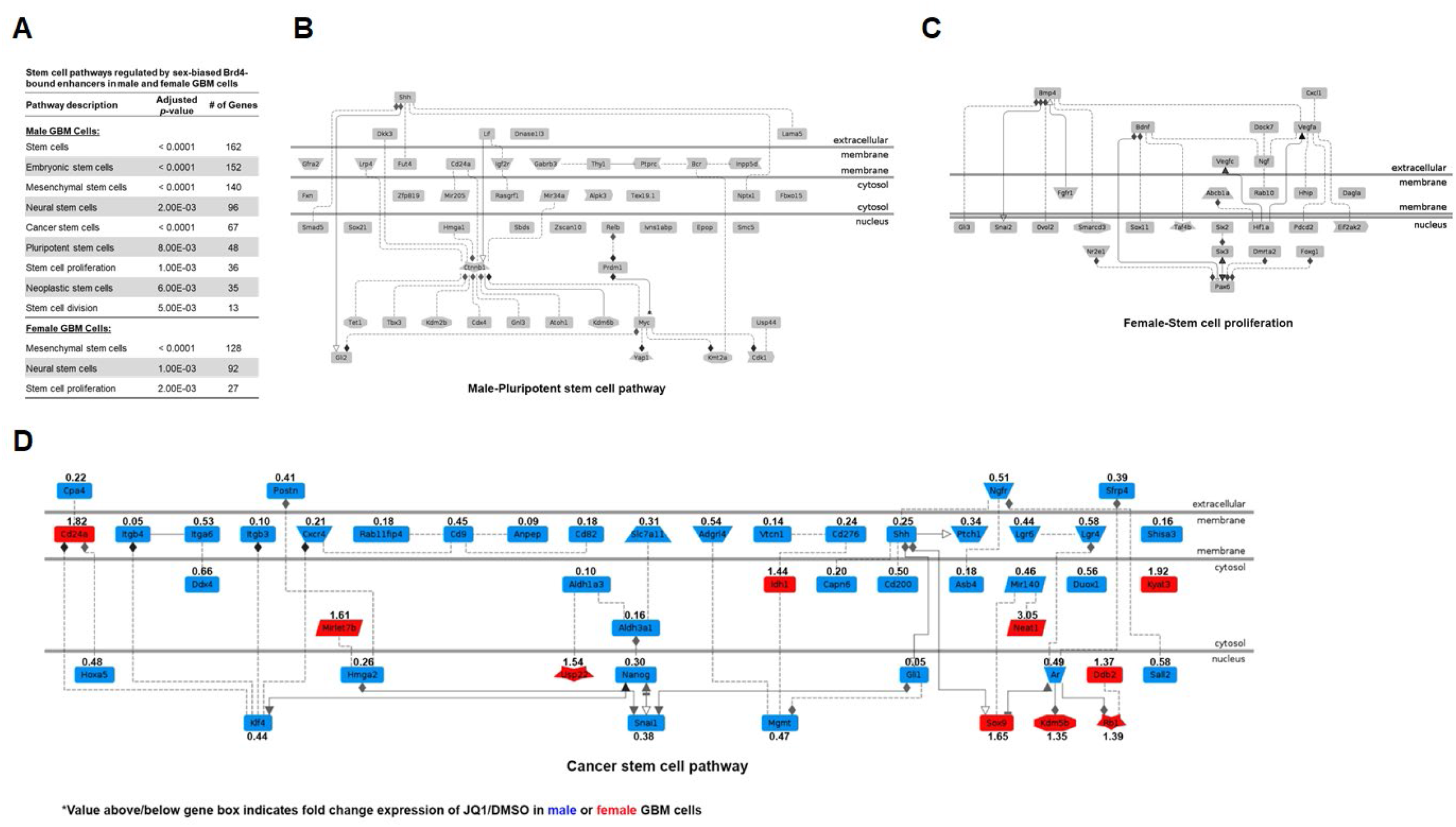
Stem cell pathways regulated by sex-biased Brd4-bound enhancers. (*A*) Pathway analysis of sex-biased Brd4-bound enhancers in male and female GBM astrocytes revealed an enrichment of stem cell pathways. Male GBM cells were highly enriched for cancer, neoplastic and pluripotent stem cell pathways. while female GBM cells were mainly enriched for neural stem cell pathways. Representative examples of these enriched pathways in male (*B*) and female (*C*) GBM astrocytes. Male-biased genes in stem cell pathways included oncogenes such as Myc, Tet1 and Lif, while female-biased genes included tumor suppressors such as Six 2 and Six 3. (*D*) Pathway analysis on sex-biased Brd4 regulated genes revealed that the cancer stem cell pathway was significantly affected by JQ1 treatment. Male genes (blue) were downregulated by JQ1 while female genes (red) were upregulated by JQ1. Value above/below gene box indicates the ratio of fold change expression of JQ1/DMSO in male (blue) or female (red) GBM cells.

### Motif analysis at sex-biased Brd4-bound enhancers identifies candidate transcription factor drivers of sex differences in the tumorigenic phenotype

Our results suggest that male and female GBM cells reside in two distinct transcriptional states established by the actions of Brd4. Inhibiting Brd4 function abrogates the sex differences in these transcriptional states and the concomitant differences in tumorigenic phenotype. As Brd4 does not bind DNA directly, but instead is recruited to enhancers by transcription factors, either directly or indirectly though histone acetylation, we sought to identify candidate transcription factors (TFs) responsible for the recruitment of Brd4 to sex-biased genomic loci. To do so, we searched for transcription factor binding motifs enriched at male-biased or female-biased Brd4-bound enhancers. Transcription factors enriched at male-biased Brd4-bound enhancers included oncogenes and stem cell markers, such as Myc, Klf5, and Oct4, whereas female-biased Brd4-bound enhancers were co-occupied and enriched with transcription factors with tumor suppressor functions, such as p53 and Smad4 (Table 1). These results could explain the observed dual functions of Brd4 in male and female GBM cells. Future studies, including knock-in and knock-out of transcriptional master regulators and partners of Brd4, are warranted to investigate the dual functions of Brd4 as an oncogene in male GBM cells and tumor suppressor in female GBM cells.

**Table 1.** Pathway analysis for enriched motifs in male and female GBM cells*

| Pathway description | Genes | Adjusted p-value | # of Genes |
| --- | --- | --- | --- |
| **<u>Male GBM Cells: 20 genes/motifs</u>** |  |  |  |
| Transcription factor activity, sequence-specific DNA binding | Myc, Hoxb4, Max, Pou5f1, Fli1, Klf1, Bach1, Fosl2, Bach2, Nfe2l2, Mafk, Isl1, Nfe2, Jun, Klf5, Fosl1, Batf, Lhx3, Atf3 | < 0.0001 | 19 |
| Neoplastic cell transformation | Myc, Hoxb4, Max, Pou5f1, Fli1, Klf1, Bach1, Fosl2, Bach2, Nfe2l2, Mafk, Nfe2, Jun, Klf5, Klf14, Fosl1, Batf, Lhx3, Atf3 | < 0.0001 | 19 |
| Cell proliferation | Myc, Hoxb4, Fosl2, Isl1, Nfe2, Jun, Fosl1, Atf3 | < 0.0001 | 8 |
| Pluripotent stem cells | Myc, Hoxb4, Max, Pou5f1, Klf1, Mafk, Isl1, Klf5, Egr1, Max, Fosl2, Jun, | < 0.0001 | 8 |
| Angiogenesis | Nfe2l2, Isl1, Jun, Klf5 | < 0.0001 | 4 |
| Mitogen activated protein kinase signaling | Max, Fosl2, Jun, Fosl1, Atf3 | 1.10E-02 | 5 |
| Stem cell differentiation | Pou5f1, Nfe2l2, Isl1, Batf, Hoxb4 | < 0.0001 | 5 |
| **<u>Female GBM Cells: 26 genes/motifs</u>** |  |  |  |
| Transcription factor activity, sequence-specific DNA binding | Mybl1, Rela, Gata4, Ets1, Atf4, Arnt, Maff, Irf1, Smad2, Irf2, Prdm1, Irf3, Ddit3, Ebf1, Smad4, Myod1, Ep300, Irf4, Cebpa, Pgr, Ctcf, Trp53 | < 0.0001 | 22 |
| Growth arrest | Mybl1, Hoxb13, Atf4, Irf1, Smad2, Ddit3, Ebf1, Myod1, Cebpa, Trp53 | < 0.0001 | 10 |
| Negative regulation of cell cycle | Mybl1, Ets1, Irf1, Ddit3, Ep300, Cebpa, Trp53 | 1.00E-03 | 7 |
| Histone modification & chromosome organization | Mybl1, Hoxb13, Atf4, Irf1, Smad2, Ddit3, Ebf1, Myod1, Cebpa, Trp53 | 1.00E-03 | 6 |
| Cyclin dependent kinase inhibitor 1 signaling | Hoxb13, Irf1, Myod1, Trp53 | 1.00E-03 | 4 |
| Estrogen receptor signaling | Hoxb13, Arnt, Irf4, Pgr | 3.00E-03 | 4 |

\* Subset of enriched pathways for enriched motifs in male and female GBM cells

### BET inhibitors have opposing effects on *in vivo* tumorigenicity in male and female mouse GBM astrocytes

The results described thus far suggested to us that Brd4-bound enhancers play an important role in maintaining sex-differences in our murine GBM model. However, these experiments all measured *in vitro* surrogates for the tumorigenic phenotype. Therefore, we next sought to determine what role, if any, Brd4 plays during *in vivo* tumorigenesis. We treated male and female murine GBM astrocytes with BETi and used these cells to perform *in vivo* tumor flank implantation studies in female nude mice. Only females were used as recipients for tumor flank implantation because we have previously shown that the sex of the recipient mouse does not affect the *in vivo* growth of implants in this murine model (19). Based on our *in vitro* ELDA studies, we chose to use JQ1 and CPI0610, the two BETi with the most dramatic sex-specific effect in male and female GBM astrocytes, respectively. Each mouse underwent flank implantation of DMSO, JQ1, or CPI0610 treated male cells (5000 cells) or DMSO, JQ1 or CPI0610 treated female cells (1.5 million cells). The sex difference in cell dose were empirically determined to match the timeframe for *in vivo* tumor growth up to the threshold for euthanasia. We attribute the sex differences in tumor growth kinetics to differences in clonogenic cell frequency and rates of proliferation (18, 19). Tumor formation was monitored by observers blinded to group assignment for 7-16 weeks. Flank implantation of JQ1-treated male GBM cells was less likely to result in tumor formation than implantation with control DMSO-treated male cells (Fig. 6 *A* and *C*). In contrast, flank implantation of CPI0610-treated female GBM cells was more likely to result in tumor formation than implantation with control DMSO-treated female cells (Fig. 6 *B* and *C*). These results are consistent with the sex-specific effects of JQ1 and CPI0610 on *in vitro* clonogenic cell frequency and cell growth. This effect was seen following only one dose of BETi prior to implantation, suggesting that this robust response is maintained and manifested at the epigenetic level. Both JQ1 and CPI0610 display a similar pattern in their selectivity to target the different Brd family members, with the highest selectivity for Brd4 (Fig. 3 *D*). Although both drugs had a similar sex-specific directional effect on clonogenic frequency in male and female cells, the magnitude of their effect differed in both the clonogenic cell frequency and *in vivo* tumorigenic assays. A possible explanation for this discrepancy is that the BETi treatment could be affecting different or multiple Brd family members. To rule out this possibility and validate that the sex-biased response to treatment was driven by Brd4, we treated male and female control and Brd4 knockdown cells with either 0.05% DMSO, 500 nM JQ1, or 1 μM CPI0610 and then performed ELDA to compare clonogenic cell frequency. No significant difference in the effect on clonogenic cell frequency was observed between control and Brd4 knockdown cells treated with BETi, indicating that Brd4 is required for sexspecific responses to BETi (*SI Appendix*, Fig. 5). Although it is still unclear why we observe different magnitude effects of these two BETi, the fact that both drugs show a sex-specific effect *in vivo* is notable and predicted by our earlier observations. These results provide strong and critical evidence that the biology of sex affects cancer incidence and outcome and addressing these sex differences in cancer will be fundamental to improving outcomes and a better quality of life for cancer patients.

**Fig. 6.**
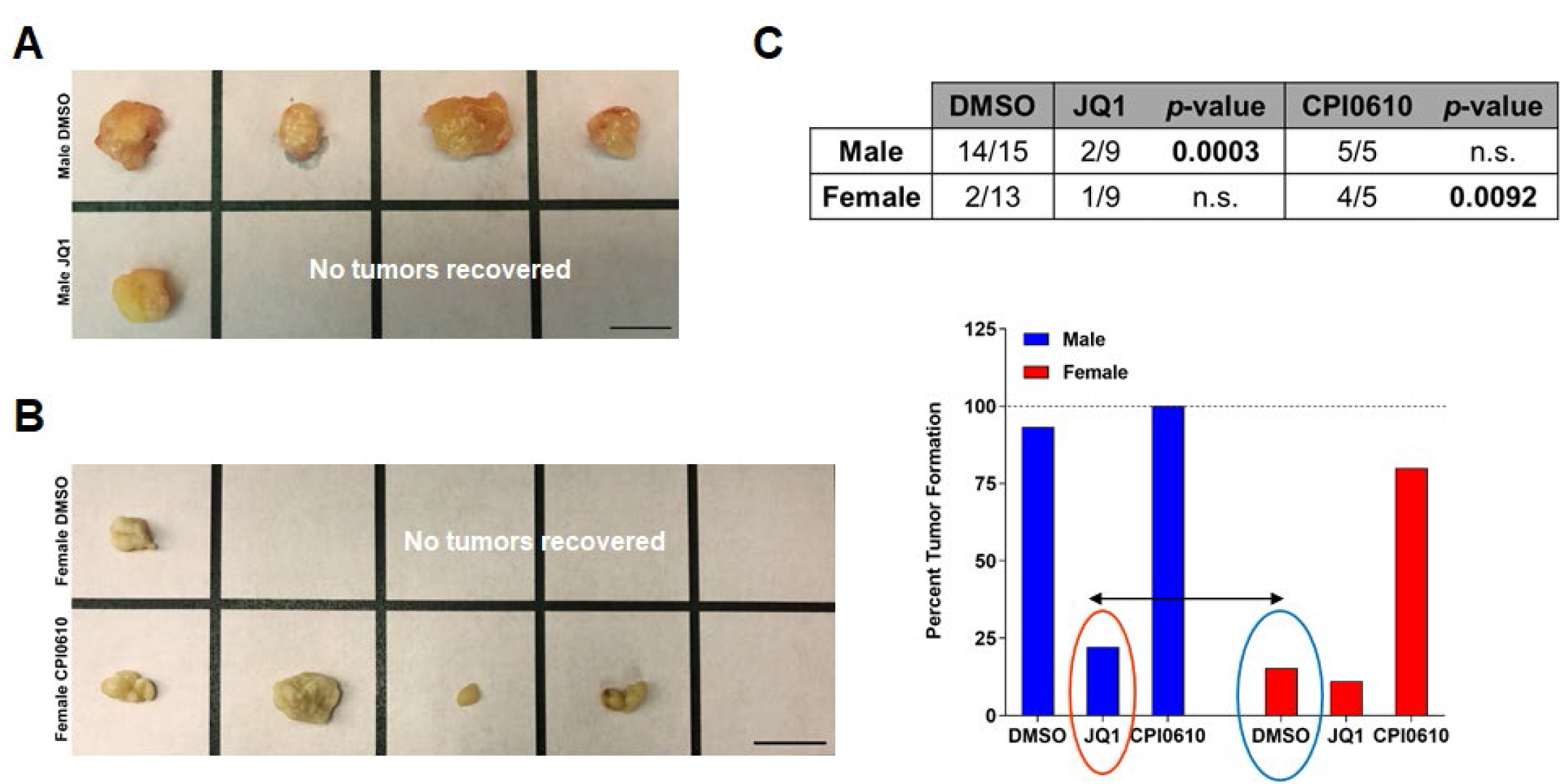
BET inhibitors have opposing effects on *in vivo* tumorigenicity in male and female GBM astrocytes. (*A*) Representative flank tumors from DMSO or JQ1 treated male GBM astrocyte-initiated tumors. (*B*) Representative flank tumors from DMSO or CPI0610 treated female GBM astrocyte-initiated tumors. (*C*) Fraction of male and female GBM cell implants that formed tumors following treatment with DMSO, JQ1 (500 nM), or CPI0610 (1 μM). JQ1-treated male GBM cells were significantly less likely to form tumors than DMSO-treated male GBM cells (Fisher’s exact test *p* <0.0003), while CPI0610-treated female GBM cells were significantly more likely to form tumors than DMSO-treated female GBM cells (Fisher’s exact test *p* <0.0092). The bar graph represents the percent tumor formation under each treatment condition. Notably, JQ1-treated male GBM cells show a decrease in their tumor formation capacity equivalent to the control DMSO-treated female GBM cells, whereas CPI0610-treated female GBM cells had an increase in tumor formation capacity almost equivalent to control DMSO-treated male GBM cells.

## Discussion

Sex differences in the incidence and severity of numerous human diseases, including cancer, are substantial and demand an understanding at a molecular level. Despite the abundant data in the literature supporting an important role for sex on incidence, prognosis, and mortality in cancer, there has been limited effort, until recently, to include sex as a biological variable in the design of clinical trials, data analysis, and treatment (3, 12, 20, 58–61). New approaches to treatment could be revealed by dissecting the biological and molecular mechanisms that drive sex differences in phenotype. Similar to other cancers, GBM has a higher incidence in males compared to females with a ratio of 1.6:1 (61). Additionally, after developing GBM, males tend to have a shorter median survival of 17.5 months compared to 20.4 months for females (62). Recently, we found that longer survival for male and female GBM patients after standard surgery, radiation, and chemotherapy is associated upon different transcriptional programs (20). Using a JIVE algorithm, male and female patients with GBM cluster into five distinct male and female subtypes distinguished by gene expression and survival. Although, a recent study by Yuan *et al*. (63) assigned GBM as a weak sex-effect cancer, their analysis of the TCGA GBM human patient data was performed using a Propensity Score Algorithm, a more standard approach compared to the specialized JIVE analysis we have previously adapted in our analysis of the TCGA GBM transcriptome datasets (20).

The epigenetic mechanisms involved in GBM initiation and treatment response have been heavily investigated in the past few years, and include epigenetic readers, writers, erasers, and histone proteins (25, 26, 64–66). The strong evidence for epigenetic dysregulation in GBM has led to clinical trials of multiple drugs targeting tumor epigenetics, in hope of improving treatment response in patients (24, 27, 29, 67). However, sex differences in the epigenetic landscape in male and female cells have not been taken into account when investigating cancer risk or treatment response. As sexual differentiation is in large part an epigenetic phenomenon and sex differences are found in the epigenetic landscape of male and female cells, it will be critical to examine the efficacy of therapies in both sexes separately to avoid missing important clinical effects when data are compiled from both sexes. Studying disparities in threshold for transformation in male and female cells will provide a powerful tool to investigate epigenetic mechanisms that establish a dimorphic tumorigenic phenotype and better treatment and survival outcomes for cancer patients.

Our study establishes that sex-biased Brd4 activity drives sex differences in GBM tumorigenic phenotype and renders male and female tumor cells differentially sensitive to BETi. Using our murine GBM model, in which male and female cells are syngeneic except in their sex chromosomes, we discovered transcriptome-wide sex differences in gene expression, H3K27ac marks, and Brd4-bound enhancers in male and female GBM cells. Brd4 is an epigenetic reader that promotes stem-like conversion and pluripotency (26, 28, 36), and its inhibition has shown therapeutic activity in a number of different cancer models (37–41). Sex-biased Brd4-bound enhancers, identified in our study, were also enriched for H3K27ac, indicative of sex-biased highly active enhancer regions. These sex-specific gene expression patterns were evident in human glioma stem-like cells, validating our model to recapitulate human GBM phenotype. Thus, for the first time, sex is established as an intrinsic element of cellular identity that is driven by Brd4 activity in GBM.

To validate whether the phenotypic differences in male and female GBM cells were indeed mediated by Brd4, as suggested by our transcriptome-wide gene expression and Brd4 binding analyses, we performed Brd4 knockdown using shRNA in male and female GBM cells. Brd4 depletion decreased clonogenic stem cell frequency in male cells, while increasing it in female cells, abrogating the sex differences observed in the tumorigenic phenotype. These results suggest that differential sex-biased Brd4 localization may be responsible for the sex differences in GBM. These data emphasize the importance of understanding and including the fundamentals of the biology of sex while investigating epigenetic mechanisms of tumorigenesis.

As bromodomain inhibitors are currently being evaluated in clinical trials, we hypothesized that this differential sex-biased Brd4 localization in male and female GBM cells would contribute to differential response to BETi. Male and female murine and human GBM cells responded in opposed ways to BETi. Consistent with our knockdown data, this dimorphism in Brd4 function was linked to the sex of the experimental cells and correlated with sex-specific patterns of Brd4 localization throughout the genome, which renders male and female cells differentially sensitive to BETi. Accordingly, BETi decreased growth of male *in vivo* tumors but increased growth of female tumors. These results strongly indicate that sex differences in disease biology translate into sex differences in therapeutic responses. The opposing sex-specific response to the inhibition of Brd4 in our GBM model could help explain previously published data in breast and prostate cancer, wherein ectopic expression of Brd4 in breast cancer cells decreased invasiveness and tumor growth, while Brd4 inhibition decreased viability of prostate cancer cells (40, 68–70). These studies also revealed that in women with estrogen receptor positive breast cancer (68) or endometrial cancer (69), low Brd4 expression correlated with worse survival. This is in contrast to men with prostate cancer, in whom low levels of Brd4 are associated with improved survival (40, 69). The sex-specific response of primary human GBM cells to BETi corroborated data obtained from the Cancer Therapeutic Response Portal that show human male GBM cells to be more sensitive to BETi, displaying slower growth and increased cell death compared to human female GBM cells.

The functions of Brd4 are determined by the action of master transcriptional regulators that bind directly or indirectly to Brd4 and dictate its localization in the genome. We performed a motif-based analysis to identify potential transcription factors that endow Brd4 with its pro- or anti-tumorigenic functions. This analysis revealed that Brd4 co-localized in a sex-biased manner with Myc and p53 in male and female GBM cells, respectively. These results are consistent with previously published data investigating sex differences in gene expression and regulatory networks in multiple human tissues. The study revealed that multiple transcription factors, although not differentially expressed between male and female cells, display sex-biased regulatory targeting patterns (2). These results warrant further functional studies to validate whether these master transcription factors drive the sex differences in Brd4-bound enhancer activity (sex-biased Brd4 genomic localization) and, therefore, the sex differences in tumorigenic phenotype and response to BETi. Given that Brd4 is shown to have pleiotropic functions in the regulation of gene expression, these results suggest that Brd4 can exhibit dual functions in GBM as an oncogene or a tumor suppressor, dependent upon cellular sex. Further functional and mechanistic experiments are warranted to ascertain this Brd4 dual function in GBM.

The consistency between our mouse and human data (GBM cell lines, GSCs and Cancer Therapeutics Response Portal) and published breast and prostate cancer studies provides strong evidence for contextdependent, sex-specific dual role of Brd4 in regulating gene expression programs in oncogenesis. In future work, we will take advantage of this remarkable sex difference in Brd4 function in GBM to better define the molecular basis for Brd4 pleiotropy in cancer and mechanisms of resistance to Brd4 inhibitors. As BETi are currently being evaluated in a number of clinical trials, understanding this phenomenon is of critical importance, both for the interpretation of existing trials and to guide better application of these drugs. Increasing our knowledge of these sex-biased genetic and epigenetic mechanisms will lead to a greater understanding of cancer biology and its relationship to normal development. We have identified sex-biased Brd4-regulated genes and pathways, which could translate into new and promising therapeutic targets to enhance survival for all GBM patients and potentially other cancers that exhibit substantial sex differences in incidence or outcome.

## Materials and Methods

### Transposon calling cards assay for Brd4 binding

For each sample (female and male murine GBM cells), 10 wells with an area of 9.5 mm^2^ were seeded at 50% confluency (200,000 cells per well of a 6-well plate). Each well represented a unique set of insertion events and was processed individually. Briefly, each well received 3 μg of plasmid DNA containing a *piggyBac* transposon (carrying a puromycin selection marker) with or without 3 μg of a transposase plasmid encoding Brd4 fused to the N-terminus of hyperactive *piggyBac* (HyPBase). Plasmids were delivered using Lipofectamine LTX/Plus Reagent (Invitrogen). Before transfection, cells were incubated in DMEM without FBS or antibiotics for 30 minutes, and then replaced with fresh media containing DMEM/F12 with 10% FBS and 1% penicillinstreptomycin. Transfection complexes were applied to cells for 12-18 hours at 37 °C and 5% CO_2_. Cells were allowed to recover in fresh medium for another 24-48 hours. Each well was then separately expanded onto 10 cm dishes and placed in medium containing puromycin at a concentration of 2.5 μg/ml for 3 days. After 3 days, cells that received transposon DNA alone showed no growth, while those that also received transposase DNA showed robust growth. Each 10 cm dish of expanded puromycin resistant cells were harvested and processed by a modified version of the transposon calling card protocol as previously described (43). The Calling Card library was prepared by assigning each well a unique Illumina P7 index, so that each well of a sample could be demultiplexed as a set of unique barcoded insertions. All 4 sets of libraries, (Male_PB_helper, Male_PB_Brd4_fusion,Female_PB_helper, and Female_PB_Brd4_fusion) were then pooled and sequenced on an Illumina HiSeq 300. A total of 40 Illumina P7 indexes were demultiplexed (10 replicates per library). Reads were mapped to the mm10 reference genome using Novoalign (version 3.08.02; Novocraft Technologies SDN BHD). For each library, data from constituent replicates were pooled and discrete insertions exported in Calling Card library format for subsequent analysis. Insertions were visualized on the WashU Epigenome Browser (v46.1).

### Sequencing data alignment and analysis

Raw reads from transposon calling cards were aligned to the murine genome build mm10 using Bowtie2 (version 2.3.4.3) (71). Significant Calling Cards peaks or Brd4 enriched enhancer sites were identified by a modified version of the previously described algorithm, which also has similarities to the MACS2 ChIP-Seq peak caller (43). Briefly, transposon insertions were grouped into peaks using a hierarchical clustering algorithm with a maximum distance of 5 Kb between insertions. Significant peaks were identified using Poisson distribution to test for enrichment over the background (unfused transposase) Calling Card data with an adjusted *p*-value threshold less than 0.05. The expected number of transposon insertions per TTAA was estimated by considering the total number of insertions observed in a large region of 100 Kb distance centered at the Calling Card cluster/insertion. We computed a *p*-value based on the expected number of insertions and identified Brd4 enriched enhancer sites using Poisson distribution.

To identify the sites with an excess of Brd4 insertions in male GBM cells relative to female cells, we used the algorithm from ChIP-Seq peaks caller MACS (version 2.1.0) (72) but modified for the analysis of calling card data. First, the Brd4-enriched enhancer sites from both male and female cells were merged if they overlapped by 1 bp. For each merged or unmerged enhancer site, the normalized insertions from the female samples were used to compute the lambda of Poisson distribution. We then computed a *p*-value from cumulative distribution function of the observed number of independent insertions in the male sample. Brd4 binding sites with *a p-*value less than 0.05 and with more than 20 insertions were considered as Brd4 binding sites significantly enriched in male sample. To identify the Brd4 sites with an excess of insertions in female GBM cells relative to male cells, we performed the same analysis, substituting the male and female data sets.

Although male and female GBM cells have a different number of X chromosomes, one X chromosome is inactivated in female cells and so the amount of accessible chromatin has been found to be similar in both cell types (73). For this reason, we treated the X chromosome in an identical manner to the autosomes for peak calling. However, to explore how sensitive our analysis was to this assumption, we recalculated *p-*values for peaks on the X chromosome as we adjusted the insertions in male cells with a multiplier from 1 to 2 (using a step size of 0.1). The total number of male-biased peaks varied from 2679 to 2995 and the total number of female-biased peaks varied from 2778 to 2727, indicating that the vast majority of differential bound peaks in our analysis are robust to the actual percentage of accessible chromatin on the inactivated X chromosome.

ChIP-seq data sets for H3K27ac were aligned to the murine genome build mm10 using Bowtie2 (version 2.3.4.3) and only uniquely aligning reads were used for downstream analyses (71). Regions of enrichment of H3K27ac over background were calculated using the MACS version (2.1.0) peak finding algorithm (72). An adjusted *p*-value threshold of enrichment of 0.01 was used for all data sets. The resulting peak files were used as inputs for DiffBind (version 3.5) to derive consensus peak sets (74, 75). The differential enrichment of H3K27Ac signals between male and female analysis was carried out with Diffbind using DESeq2 (method = DBA_DESEQ2) with libraries normalized to total library size.

RNA-seq data sets were aligned to the transcriptome and the whole-genome with STAR (version 2.7.0) (76). Genes or exons were filtered for just those that were expressed. The readcount tables for annotated genes in the mm10 gene transfer format (.GTF) file were derived from uniquely aligned reads using HTSeq (version 0.11.1) (77). The raw counts for each gene were converted into TPM that is more appropriate for comparisons of gene expression levels between samples (78–80). TPMs from biological replicates were averaged for subsequent analysis. Differential gene expression between pairs of samples was computed using DESeq2 (version 1.28.1) and was filtered by FDR < 0.05 for differentially expressed genes (81). ln in some cases, a two-fold change was also applied as a filter for identifying differentially expressed genes between two conditions.

### Data Availability

Please find in Sl Appendix, Supplementary Materials and Methods detailed description of Mouse male and female GBM cells, Chromatin immunoprecipitation sequencing (ChIP-seq) for H3K27ac, RNA-sequencing, Pathway analysis, Human glioma stem-like cells (GSCs) gene expression data analysis, Quantitative Real-Time PCR, shRNA lentiviral infection and knockdown of Brd2, Brd3 and Brd4, Clonogenic cell frequency assay: ELDA analysis, Growth assays, Cancer Therapeutic Response Portal data analysis, *In vivo* tumorigenesis: flank implantation, Statistical analysis and Bioinformatic motif analysis. All Illumina sequencing reads and processed files are publicly available via an FTP server in the Center for High-Performance Computing at Washington University. The server ftp links are below: Calling Card data and processed files: https://htcf.wustl.edu/files/pX76jgeW ChlP-seq data and processed files: https://htcf.wustl.edu/files/LdbbjVd3 RNA-seq data and processed files: https://htcf.wustl.edu/files/NeAbNQX2 Additionally, these files have been deposited in the Short Read Archive/GEO database (http://www.ncbi.nlm.nih.gov/geo/). GEO accession # in process.

## ACKNOWLEDGMENTS

This work was supported by NIH RO1 CA174737 (JBR), R01 NS076993 (RM), R21 HG009750 (RM), U01 MH109133 (RM), RF1 MH117070 (RM), Joshua’s Great Things (JBR), The Children’s Discovery Institute (JBR and RM) and NIH T32GM007200, T32HG000045, and F30HG009986 (AM). We thank the Genome Technology Access Center in the Department of Genetics at Washington University School of Medicine for help with genomic analysis. The Center is partially supported by NCI Cancer Center Support Grant #P30 CA91842 to the Siteman Cancer Center and by ICTS/CTSA Grant# UL1 TR000448 from the National Center for Research Resources (NCRR), a component of the National Institutes of Health (NIH), and NIH Roadmap for Medical Research. This publication is solely the responsibility of the authors and does not necessarily represent the official view of NCRR or NIH.

## Author contributions

NK and ZQ designed and performed experiments, analyzed data and wrote the manuscript. BP performed all the bioinformatic analyses for the glioma stem-like cells data. MW and AM developed and optimized the calling cards protocol. MW and XC performed the calling card experiments and generated libraries. LB provided significant edits to the manuscript and suggestions to the experiments and data analyses as well as blindly analyzed ELDA experiments. AM developed the software to analyze and visualize calling card data. KB and JG performed ChIP-seq experiments. SS assisted in the RNA-seq analysis. JNR provided significant edits to the manuscript and suggestions for human samples experiments. JBR and RM designed experiments, analyzed data, edited the manuscript and supervised the project. All authors read and approved the final version of the manuscript.

## Supplementary Information for

### Supplementary Materials and Methods

#### Mouse male and female GBM cells

Male and female astrocytes were isolated from the neocortices of postnatal day 1 Nf1fl/fl GFAP Cre mice and genotyped for sex using Jarid1c and Jarid1d PCR. Male and female Nf1−/− astrocytes were then infected with retrovirus encoding a flag-tagged dominant-negative form of p53 (DNp53) and EGFP resulting in male and female astrocytes null for Nf1 and p53 function. The DNp53 plasmid consists of amino acids 1–14 of the transactivation domain followed by amino acids 303–393, thus lacking the DNA binding domain. Cells were grown in DMEM/F12 media supplemented with 10% FBS and 1% penicillinstreptomycin. These astrocytes serve as a model of GBM and are referred to throughout the text as male and female GBM astrocytes. Cell cycle analysis and metaphase spreads of male and female GBM astrocytes reveal similar aneuploidy with 2N and 4N subpopulations with varying and overlapping chromosome numbers that ranged between 50 and 120, and 41 and 111 respectively (1).

#### Chromatin immunoprecipitation sequencing (ChIP-seq) for H3K27ac

10 million male and female GBM cells were treated with 0.05% DMSO or 500 nM JQ1 for 24 hours. Following treatment, cells were then prepared for various downstream analyses including ChIP-seq for H3K27ac. Briefly, cells were fixed in 1% formaldehyde prepared in DMEM/F12 and incubated for 10 minutes at room temperature with gentle rocking. Fixation was quenched by the addition of glycine to a final concentration of 125 mM and incubation for 10 minutes at room temperature with gentle rocking. Cells were then washed with cold PBS and harvested by scraping in 5 ml of cold PBS. Cells were transferred to 15 mL conical tubes and pelleted with a 5-minute spin (1,000g) at 4°C. Sonication was performed on an Epishear probe-in sonicator (Active Motif) at 40% amplitude for 8 cycles of 30 seconds, with 30 seconds of rest between each cycle. ChIP-seq libraries were constructed as previously described (2), with an antibody that recognizes H3K27Ac (Active Motif, 39133, rabbit polyclonal). ChIP-seq libraries were constructed with sample-specific barcodes, and pooled before sequencing on a HiSeq 2500 (Illumina).

#### RNA-sequencing

Male and female GBM cells (Nf1−/−;DNp53 astrocytes) were generated as previously reported (Sun et al., 2015) and grown in DMEM/F12 media supplemented with 10% FBS and 1% penicillin-streptomycin. Total RNA was isolated from male and female GBM cells that were treated with DMSO (0.05%) or JQ1 (500 nM for 24 hours) using the RNeasy Mini Kit from QIAGEN (Hilden, Germany), following the kit protocol. PolyA Selection was performed to create RNA Seq libraries. Cell mRNA was extracted from total RNA using a Dynal mRNA Direct kit. The quantity of RNA was measured using a spectrophotometer (NanoDrop 2000c; Thermo Scientific). Samples with an RNA concentration A260/A280 ≥1.8 ng/μl, and purity A230/A260 ≥ 2.0 ng/μl were selected. The Agilent 2100 Bioanalyzer was used to determine the RNA integrity number. The degradation level was identified using the RNA 6000 Nano LabChip kit (Agilent). Samples with RNA integrity number > 9.8 were further processed using TruSeq mRNA Library Preparation Kit (Illumina) and then sequenced on a HiSeq 25000 (Illumina).

#### Pathway analysis

Pathway enrichment analysis for differentially regulated genes was performed using a combination of KEGG pathway and Genomatix Pathway System (GePS). GePS uses information extracted from public and proprietary databases to display canonical pathways and or to create and extend networks based on literature data. These sources include NCI-Nature Pathway Interaction Database, Biocarta, Reactome, Cancer Cell Map, and the ENCODE Transcription Factor project data. All data for pathway analyses are presented with adjusted corrected *p*-values.

#### Human glioma stem-like cells (GSCs) gene expression data analysis

RNA-sequencing data were derived from GSCs published in Mack et al., 2019 (3). Data from FASTQ files were trimmed with TrimGalore, which implements Cutadapt (4) and FastQC (5), to remove low quality reads and trim adapaters. Reads were aligned to Gencode v29 using Salmon (6) with correction for GC bias, positional bias and sequence-specific bias. The R/Bioconductor (7) package tximport (8) was used to generate TPM values. Low expressed genes without at least 1 count in 4 samples were excluded from analysis. Comparisons were performed using DESeq2 (9) on raw counts.

#### Quantitative Real-Time PCR

Total RNA was isolated using QIAGEN RNeasy Mini Kit from male and female GBM cells (Nf1−/−;DNp53 astrocytes) infected with shRNA lentivirus against Brd2, Brd3 or Brd4. cDNA was generated using the QuantiTect Reverse Transcription Kit (Qiagen). Quantitative RT-PCR was performed using gene-specific primers and iTaq SYBR Green PCR master mix (Biorad, CA). Data was analyzed by standard ΔCq method (2-ΔΔCq) where ΔCq is the difference between the gene of interest and GAPDH control Cq value. Sequences for primers specific for each Brd family member are provided in *SI Appendix,* Table 5.

#### shRNA lentiviral infection and knockdown of Brd2, Brd3 and Brd4

Brd2, Brd3 and Brd4 knockdown lines were generated by infecting male and female GBM cells (Nf1−/−;DNp53 astrocytes) with lentiviral shRNAs against Brd2, Brd3 or Brd4. All Knockdown lines were selected with puromycin (2.5 μg/ml) in media for 1-2 weeks and the survivors were expanded for downstream target knockout analysis. Five different shRNAs specific to each of the Brd family members were evaluated and the shRNA with the most robust knockdown was used for the downstream functional experiments. Sequences for shRNAs are provided in *SI Appendix,* Table 5.

#### Clonogenic cell frequency assay: Extreme Limiting Dilution Assays (ELDA analysis)

Male and female mouse GBM cells (Nf1−/−;DNp53 astrocytes, 2 lots of male and female cells) or human male and female primary GBM cell lines (3 male and 3 female human primary GBM cell lines were kindly provided to us by Dr. Albert Kim) were treated with DMSO (0.05%), iBET762 (4 μM), CPI203 (0.5 μM), CPI0610 (1 μM), JQ1 (500 nM), OTX015 (2.5 μM), PFI1 (5 μM) or RVX-208 (5 μM) for 24 hours or with shRNAs against Brd2, Brd3, or Brd4. The timing and dosing of each drug was based on previously published data (10–14). Clonogenic capacity was then assessed by the Extreme Limiting Dilution Assay (ELDA). The frequency of clonogenic stem cells was evaluated by the cells’ ability to form tumor-spheres in low-adherent conditions as previously reported (15). Briefly, cells were harvested into a single cell suspension and plated in neurosphere media containing EGF, FGF and mouse LIF on 96-well ultra-low attachment plates in a serial dilution ranging from 3000 cells/well to 1 cell/well (3000, 600, 120, 24, 5 and 1 cells; n=14-18 wells/cell density). Sphere formation was measured 7 days after plating. Clonogenic cell frequency was then analyzed using the Extreme Limiting Dilution Analysis software (16).

#### Growth assays

Growth kinetics of male and female GBM cells (Nf−/−;DNp53 astrocytes) treated with DMSO (0.05%), JQ1 (500 nM), or RVX-208 (5μM) for 24 hours were examined by counting live cell number using an automated T4 cell counter as previously described with minor modifications (15). Briefly, cells were harvested and plated in a 6-well plate at a density of 2 x 10^4^ cells/well (2 technical replicates per treatment/genotype/time point). 4 hours post plating, cells were harvested by trypsinization and counted in the presence of trypan blue. This time point was designated as the starting point (T0) of the time course. Cells were then harvested and counted every 24 hours for a total of 3 days (24, 48 and 72 hours). This experiment was repeated three times.

#### Cancer Therapeutic Response Portal data analysis

Drug response data were derived from the Cancer Therapeutic Response Portal (CTRP). The outcome variable was area under the curve (AUC) for each cell line and drug treatment. Glioblastoma cell lines were segregated by male vs. female sex, and AUC for each drug was analyzed by sex using the R/Bioconductor package limma (17).

#### *In vivo* tumorigenesis: flank implantation

Flank tumors were generated by implanting GBM astrocytes subcutaneously into left side flanks. These cells were treated with 0.05% DMSO, 500 nM JQ1 or 1 μM CPI0610 for 24 hours, followed by EGF treatment at 50ng/ml for one week. 1.5 million female cells or 5000 male cells treated with DMSO, JQ1 or CPI0610 were then harvested and resuspended in 100 μl of 1:1 media to matrigel (BD Biosciences) and injected into the flanks of mice. Mice were monitored weekly and tumor growth and formation were monitored blindly for 8-16 weeks with thrice weekly micrometer measurements in 3 dimensions. Animals were used in accordance with an animal studies protocol (no. 20180206) approved by the Animal Studies Committee of the Washington University School of Medicine per the recommendations of the Guide for the Care and Use of Laboratory Animals (NIH).

#### Bioinformatic motif analysis

To identify DNA motifs that are associated with sex-biased Brd4 binding, we used the HOMER toolkit (v4.11) (18). Briefly, to find motifs enriched under male-biased peaks, we used the sequences under male-biased Brd4 peaks as input to HOMER, and used sequences from female-biased peaks, or all Brd4 peaks, as background. We used the following HOMER command: (findMotifsGenome.pl mm10 -nomotif -preparsedDir -bg). We found motifs enriched under female-biased peaks in a similar fashion using the corresponding sequence files.

#### Statistical analysis

All experiments in this study were carried out at least three times. ELDA experiments with JQ1 and CPI0610 treatment were performed three times (n=3) in two or three different mouse GBM lots of cells, as indicated for each experiment. ANOVA, two-tailed Student’s t-test and Fisher’s exact test were used to compare the differences in all functional measurements between BET inhibitors and control group (DMSO), and a p-value < 0.05 was considered statistically significant.

To test if upregulated genes in male cells compared to female cells and vice versa are significantly enriched for H3K27ac binding, the normalized H3K27ac signal intensities (reads) from 1kb upstream of the gene start site to 1kb downstream of the end of the gene were used for the correlation analysis. A paired Mann-Whitney-Wilcoxon test was used to compare normalized H3K27ac signal intensities between male and female cells and a p-value less than 0.01 was considered statistically significant.

To investigate if Brd4-proximal genes are significantly downregulated upon JQ1 treatment compared to Brd4-distal genes, we first defined Brd4 proximal genes as the closest genes to Brd4 binding sites and Brd4 distal genes as genes located near sites that are not enriched for Brd4 binding sites. Brd4 proximal and distal genes (500 genes each) were randomly selected. A Mann-Whitney-Wilcoxon test was used to compare the expression profiles before and after JQ1 treatment of Brd4 proximal and distal genes for male and female respectively and a p-value less than 0.01 was considered statistically significant.

**Fig. S1.**
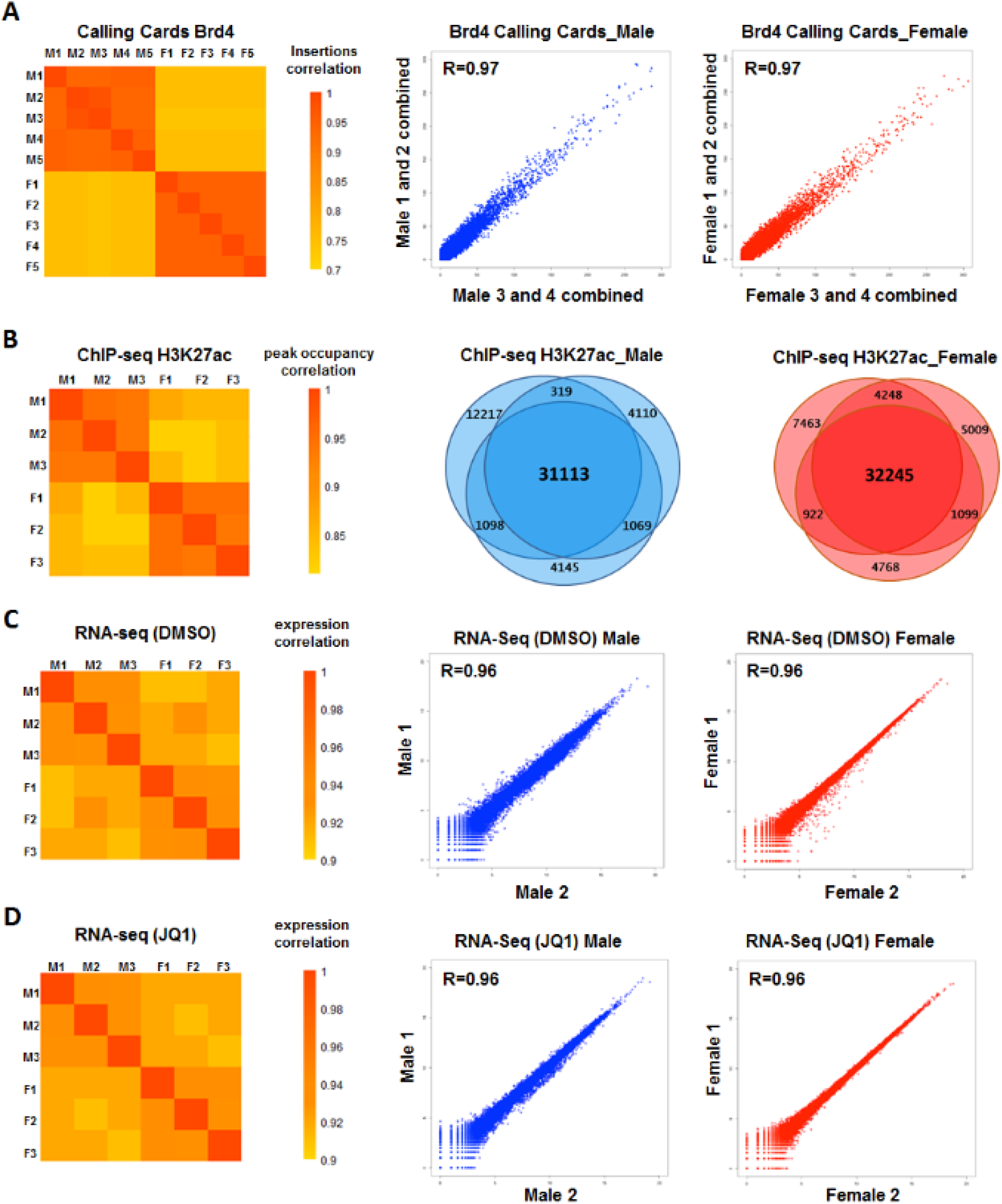
Quality assessment of transposon Calling Card, H3K27ac ChIP-seq and RNA-seq experiments. Correlation heatmaps for transposon Calling Card *(A)*, H3K27ac ChIP-seq *(B)* and RNA-seq *(C, D)* were prepared using replicates of male and female GBM samples. Venn diagrams in *(B)* depict the shared H3K27ac-enriched peaks among replicates in male and female GBM cells.

**Fig. S2.**
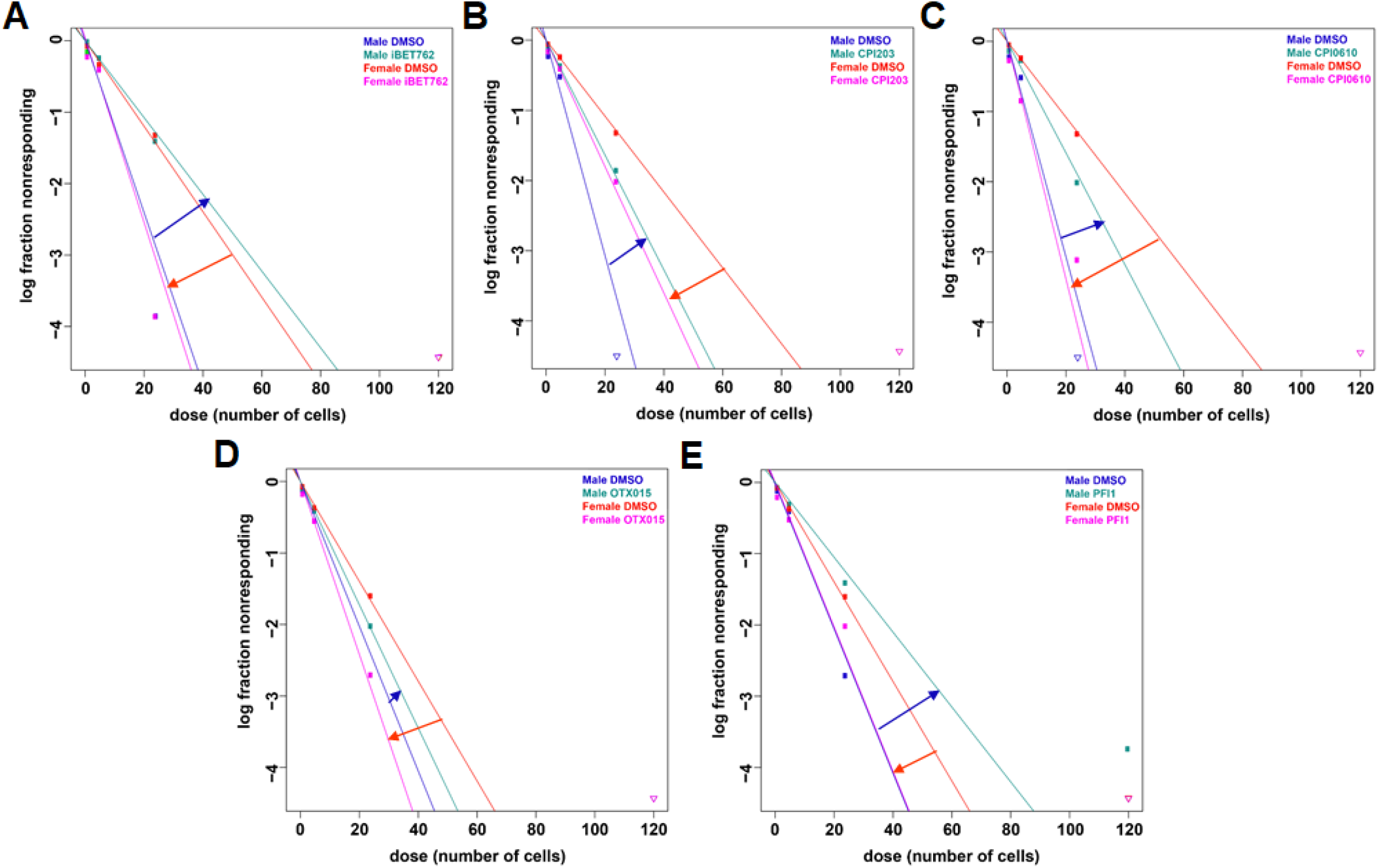
BET inhibitors have opposing effects on clonogenic stem cell frequency in male and female GBM cells. Frequency of clonogenic stem-like cells as determined by ELDA assay in male and female GBM cells treated with DMSO or *(A)* iBET762, *(B)* CPI203 *(C)* CPI0610 *(D)* OTX015 *(E)* PFI1. Male cells exhibited greater clonogenic cell activity than female cells under control conditions. BET inhibition reduced clonogenic cell frequency in male cells, while female cells exhibited an increase in their clonogenic cell frequency.

**Fig. S3.**
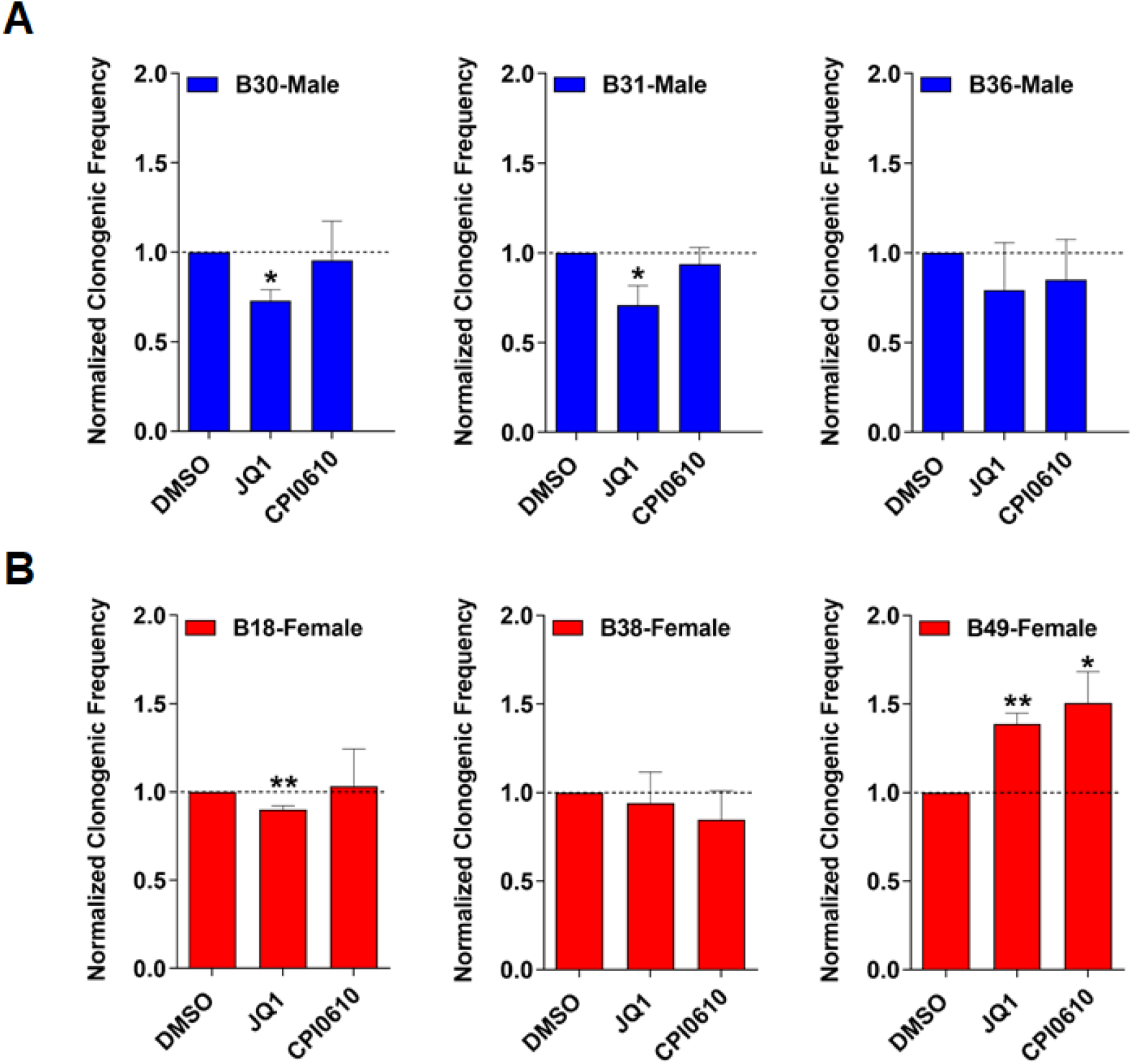
Sex-specific response to BET inhibitor treatment is evident in human primary GBM cells. Normalized clonogenic cell frequency as determined by ELDA assay in male *(A)* and female *(B)* human primary GBM cells treated with DMSO (control), JQ1, or CPI0610. JQ1 treatment significantly reduced clonogenic cell frequency in human primary GBM male cells, and significantly increased clonogenic cell frequency in 1 out of 3 human primary GBM female cells. CPI0610 treatment had a similar response pattern as JQ1 in male and female human GBM primary cells. (*=*p* <0.05, **=*p* <0.01 as determined by unpaired two-tailed t-test).

**Fig. S4.**
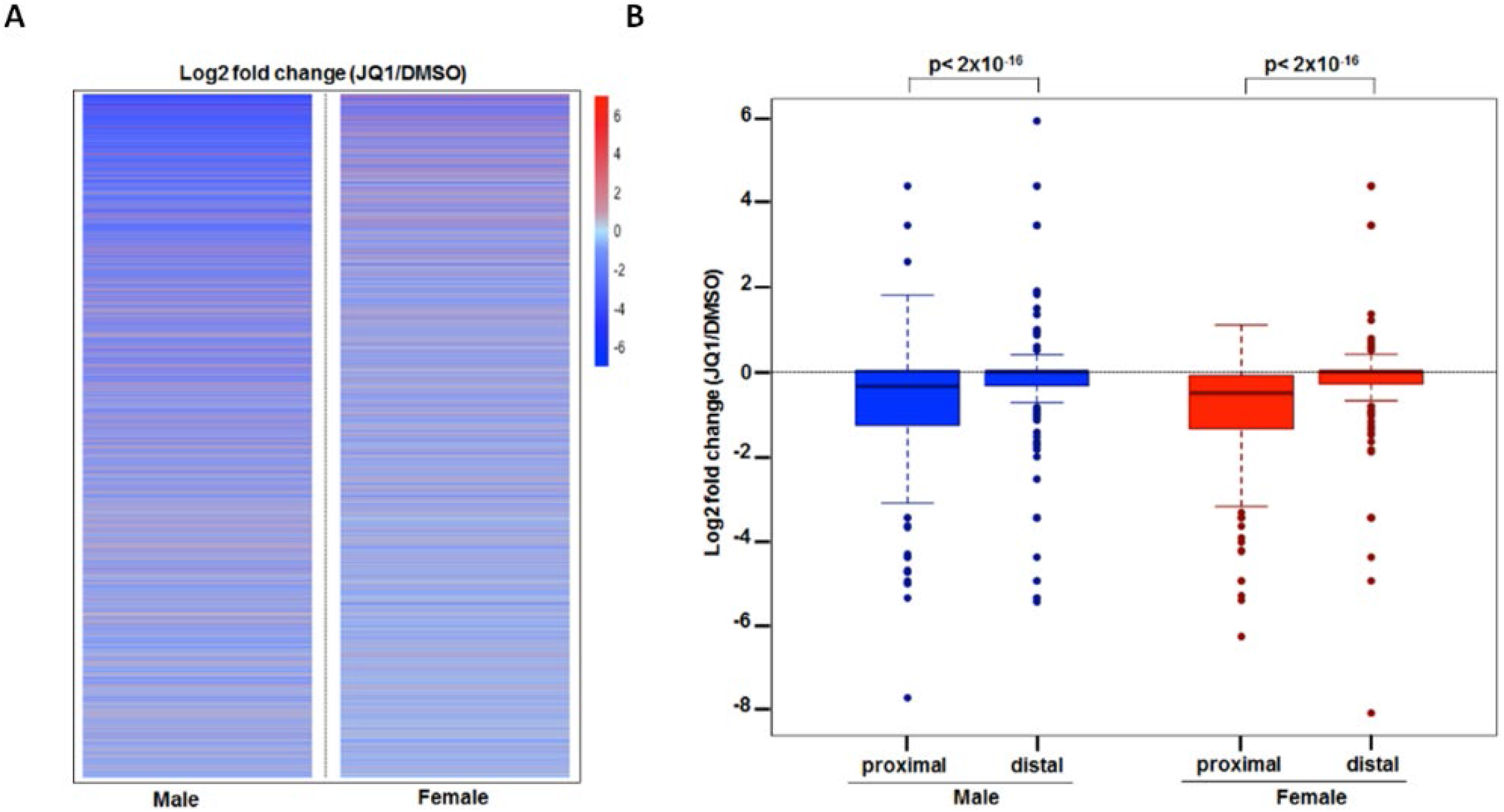
JQ1 effect on gene expression in male and female GBM cells. *(A)* Heat map of gene expression changes (log2; upregulated in red and downregulated in blue) upon JQ1 treatment in male and female GBM cells. *(B)* Boxplot of gene expression changes (log2) of Brd4 proximal and distal genes following JQ1 treatment of male (blue) and female (red) GBM cells. The gene expression profile of genes in close proximity to Brd4 binding sites is compared to distal genes by a Mann-Whitney-Wilcoxon test. Brd4 proximal genes are significantly downregulated compared to Brd4 distal genes in both male and female GBM cells following JQ1 treatment (*p*< 2×10^-16^).

**Fig. S5.**
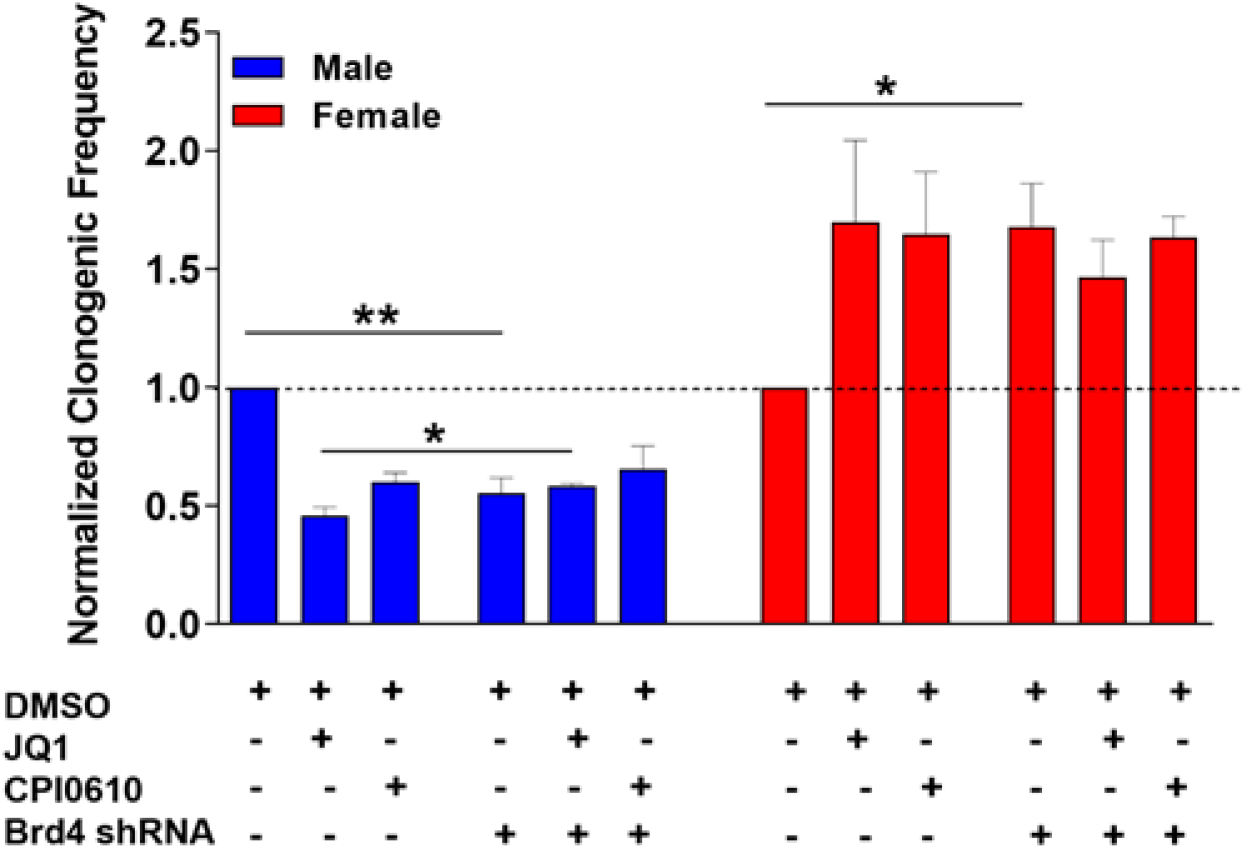
Brd4 is the main Brd family member responsible for the sex-specific response to BETi treatment. Normalized clonogenic cell frequency as determined by ELDA assay in male and female GBM cells treated with DMSO or the indicated BET inhibitor, with or without Brd4 knockdown. No significant difference in the effect on clonogenic cell frequency is observed between control and Brd4 knockdown cells treated with BET inhibitors, indicating that Brd4 is the main Brd family member responsible for this sexually dimorphic response to BETi treatment in male and female cells. (*=*p* <0.05, **=*p* <0.01 as determined by unpaired two-tailed t-test).

**Table S1.** Pathway enrichment analysis for sex-biased Brd4-regulated genes in male and female GBM cells as identified by Brd4 calling cards and H3K27ac ChIP_seq*

| Pathway description | Adjusted *p*-value | # of Genes |
| --- | --- | --- |
| **<u>Male GBM Cells: 1740 genes</u>** |  |  |
| Glioblastoma | < 0.0001 | 349 |
| Cell proliferation | < 0.0001 | 224 |
| Cell cycle | < 0.0001 | 152 |
| Regulation of transcription | < 0.0001 | 113 |
| Tumor angiogenesis | 5.00E-03 | 70 |
| Cancer stem cells | < 0.0001 | 41 |
| Epidermal growth factor receptor signaling | 2.00E-03 | 15 |
| **<u>Female GBM Cells: 1604 genes</u>** |  |  |
| Cell differentiation | < 0.0001 | 386 |
| Glioblastoma | < 0.0001 | 320 |
| Regulation of transcription | < 0.0001 | 181 |
| Cell migration | < 0.0001 | 161 |
| Neural stem cells | < 0.0001 | 72 |
| Negative regulation of angiogenesis | 1.00E-03 | 13 |
| Kinase suppressor of Ras | 1.00E-03 | 8 |

\*Subset of pathway enrichment analysis

**Table S2.**
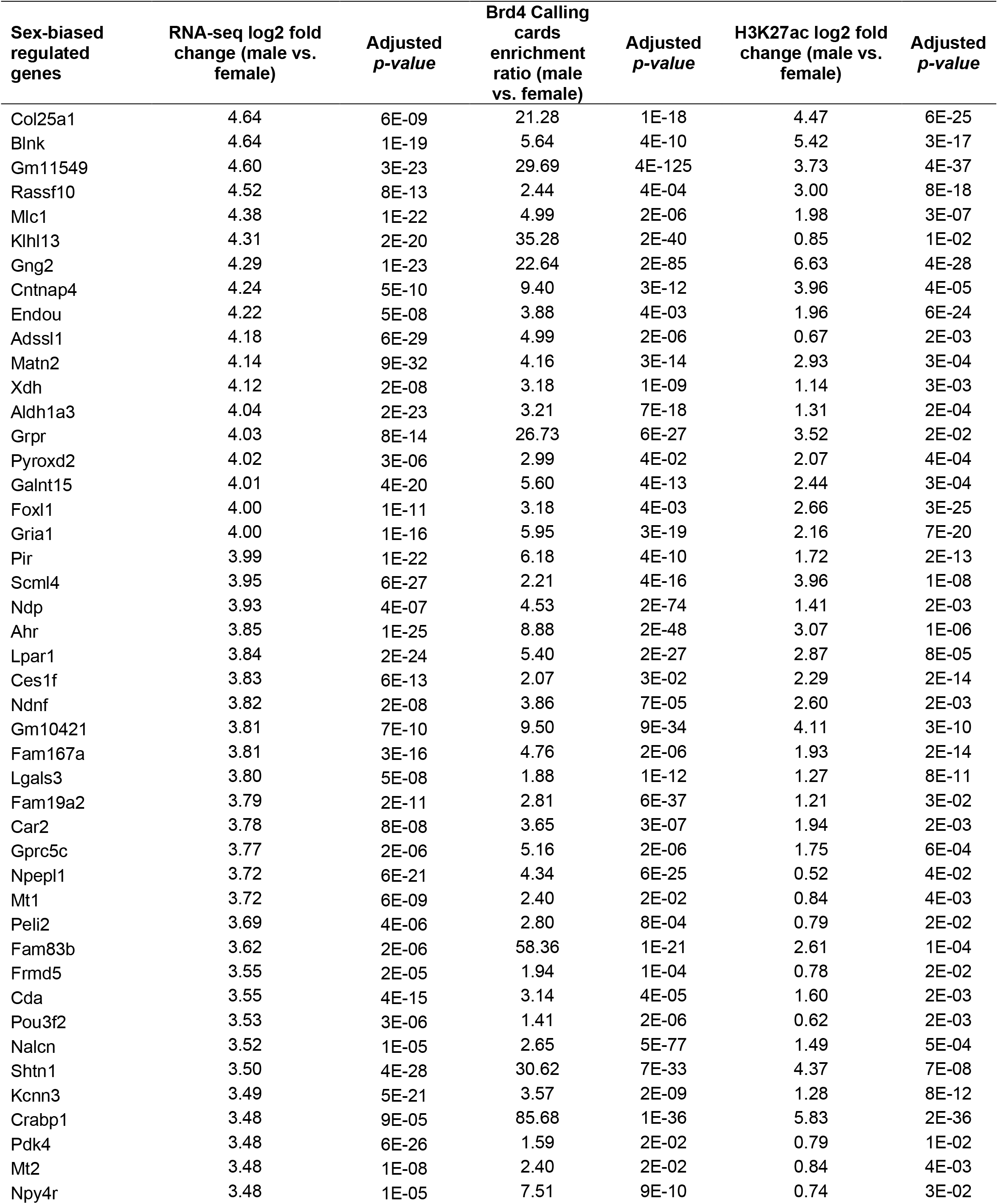
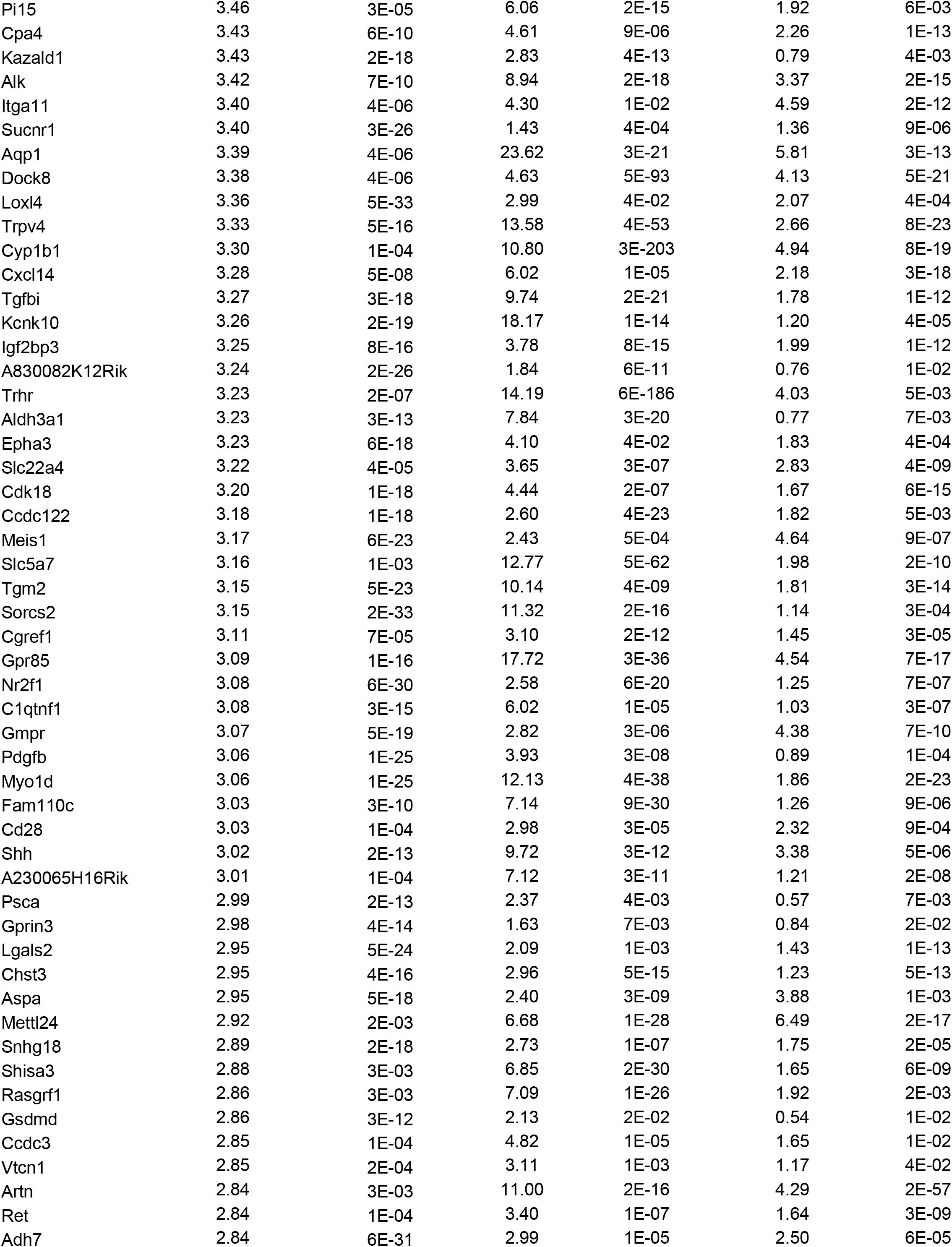
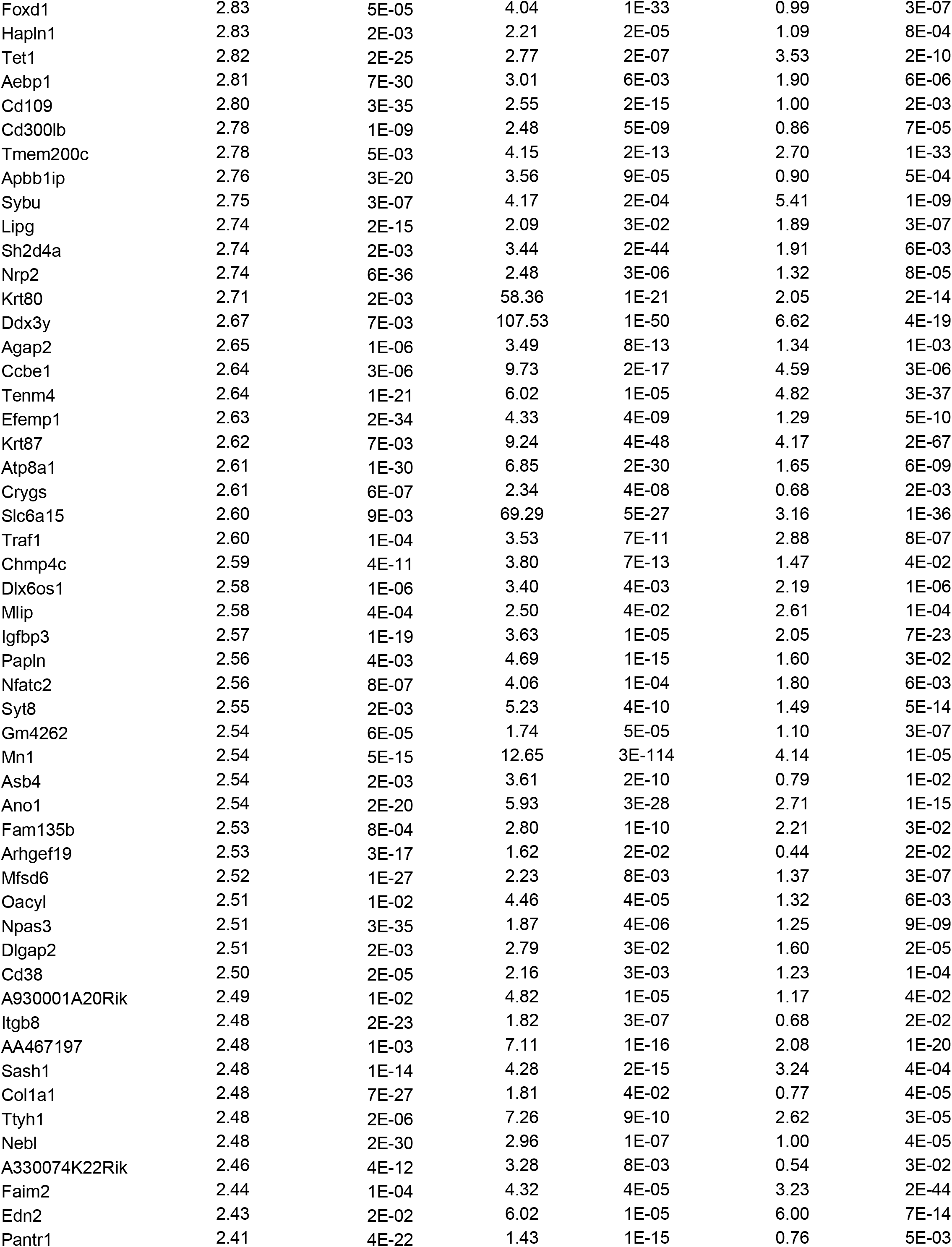
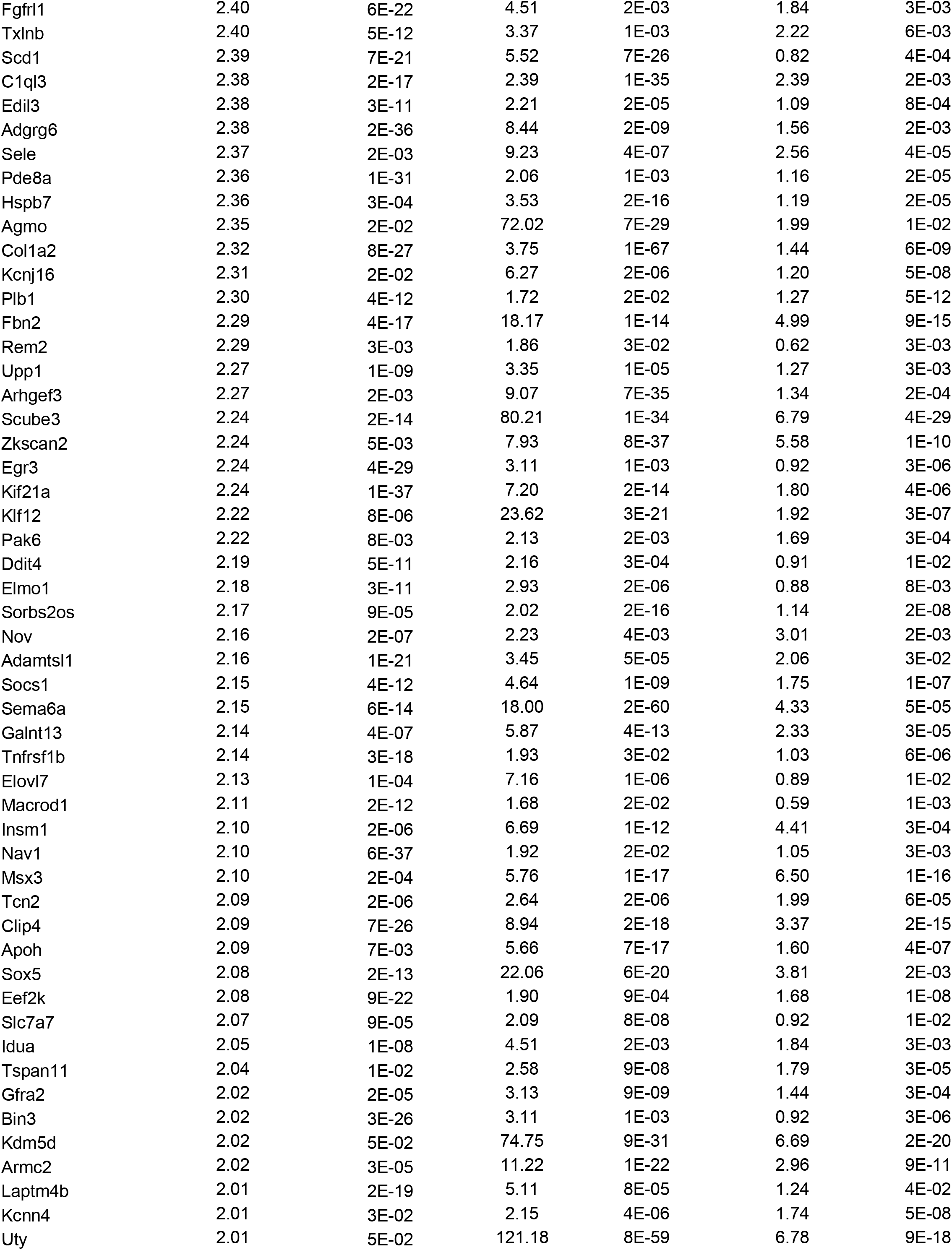
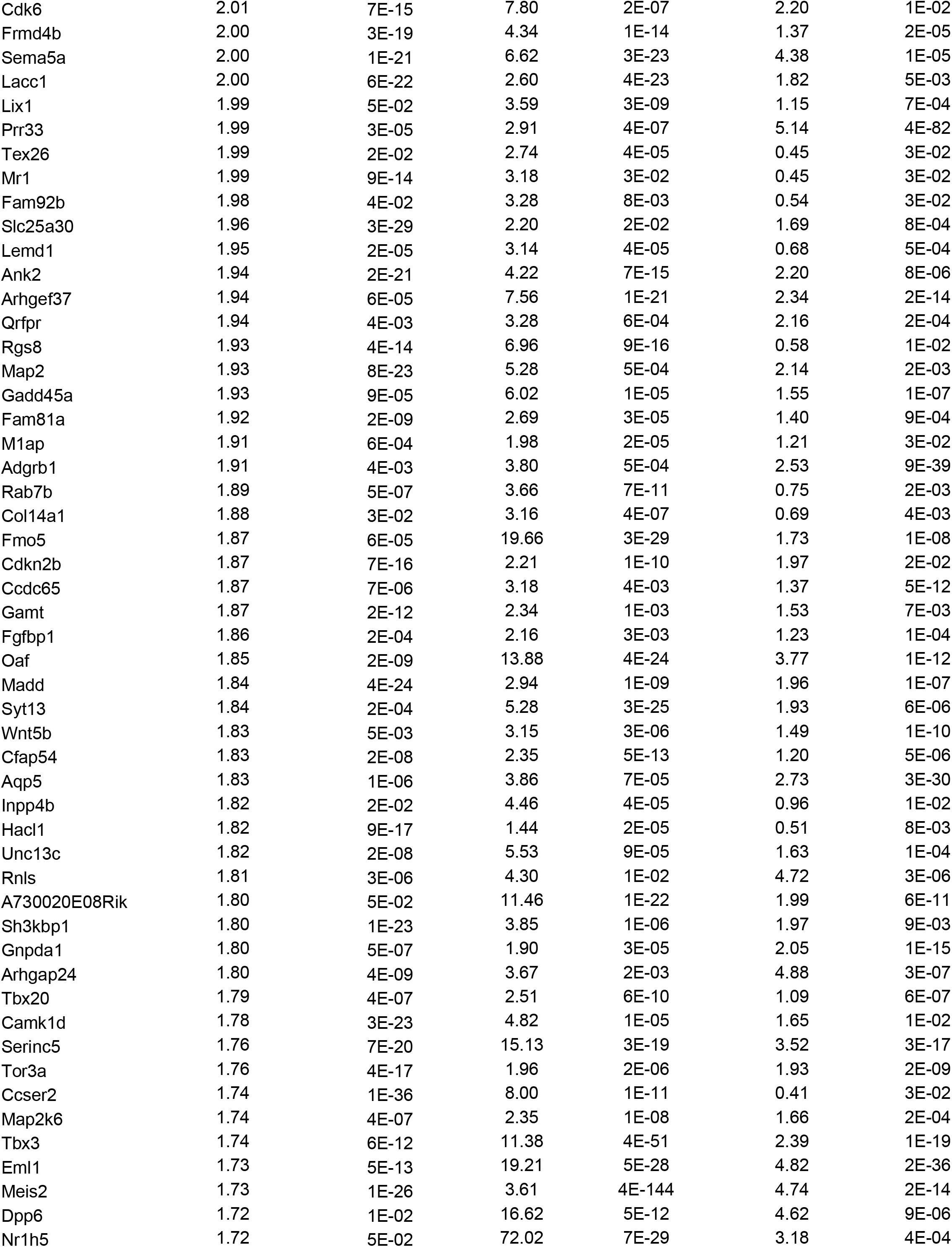
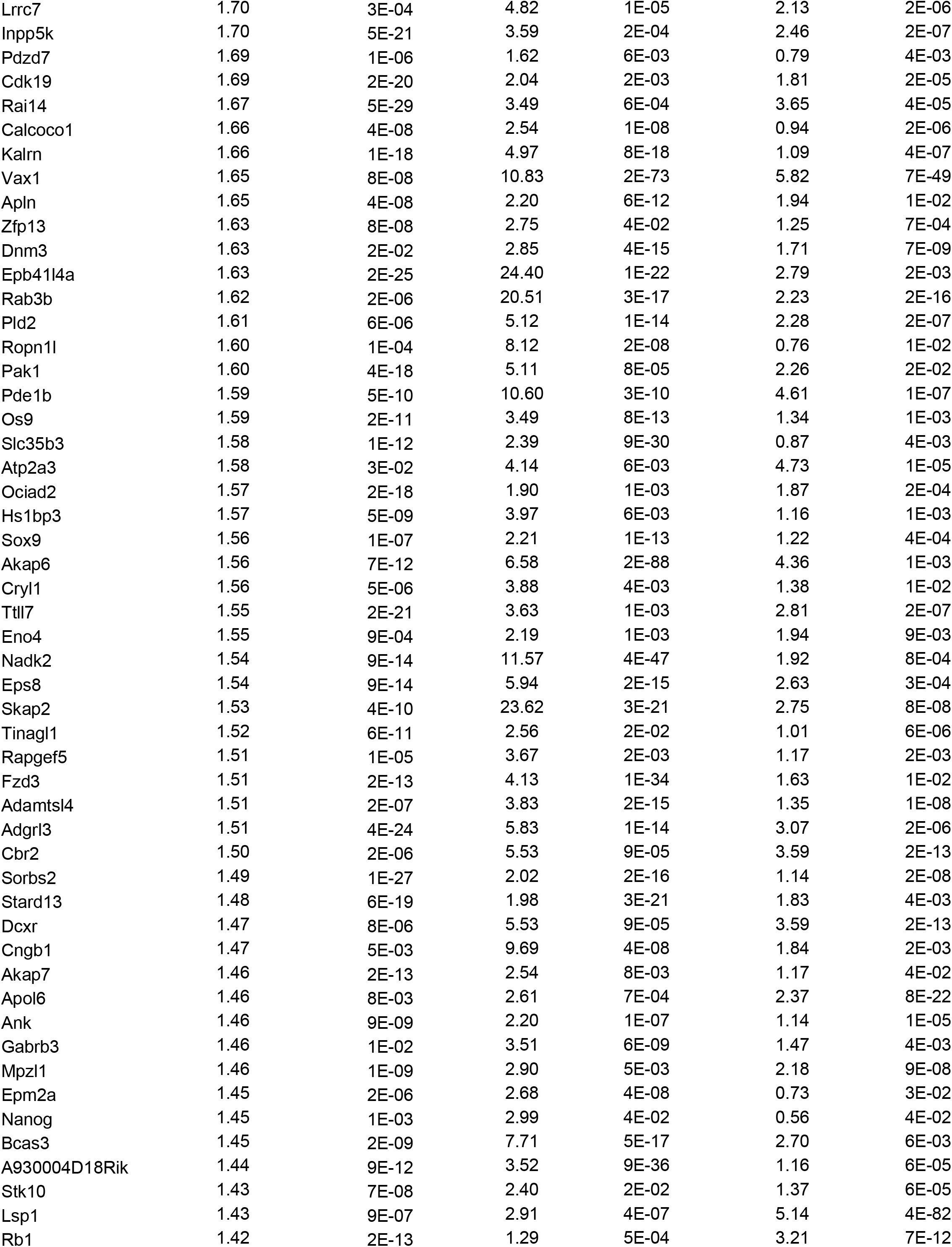
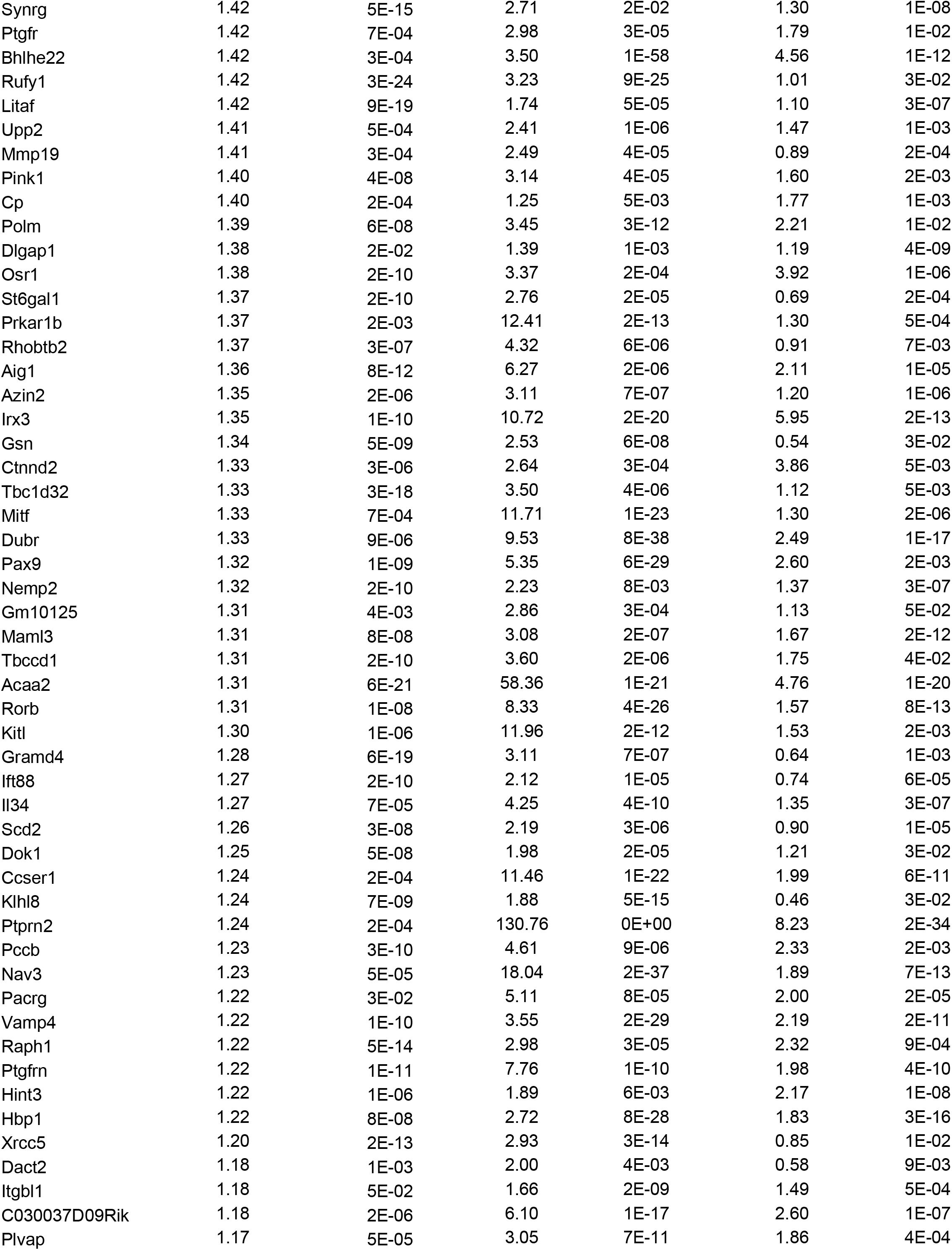
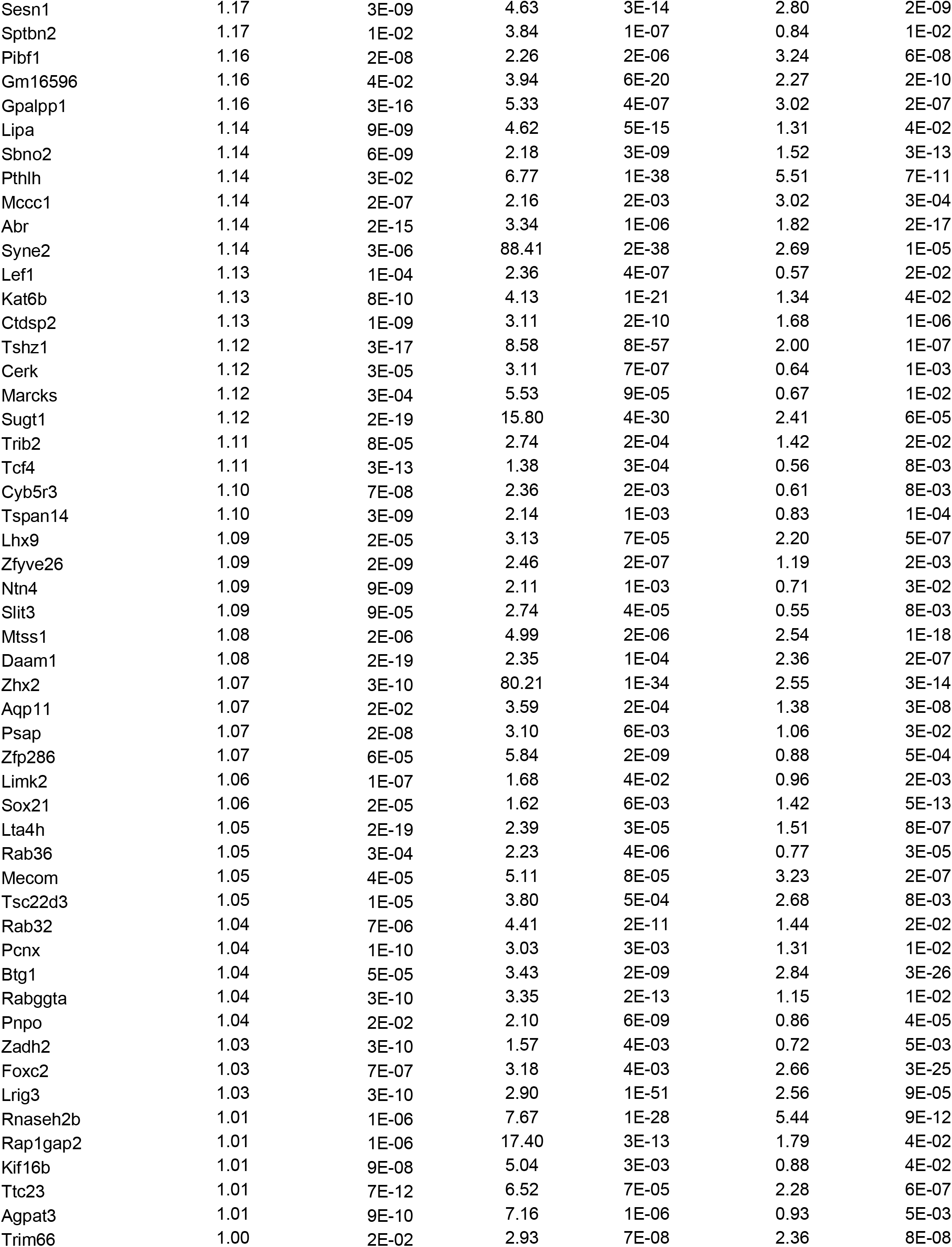
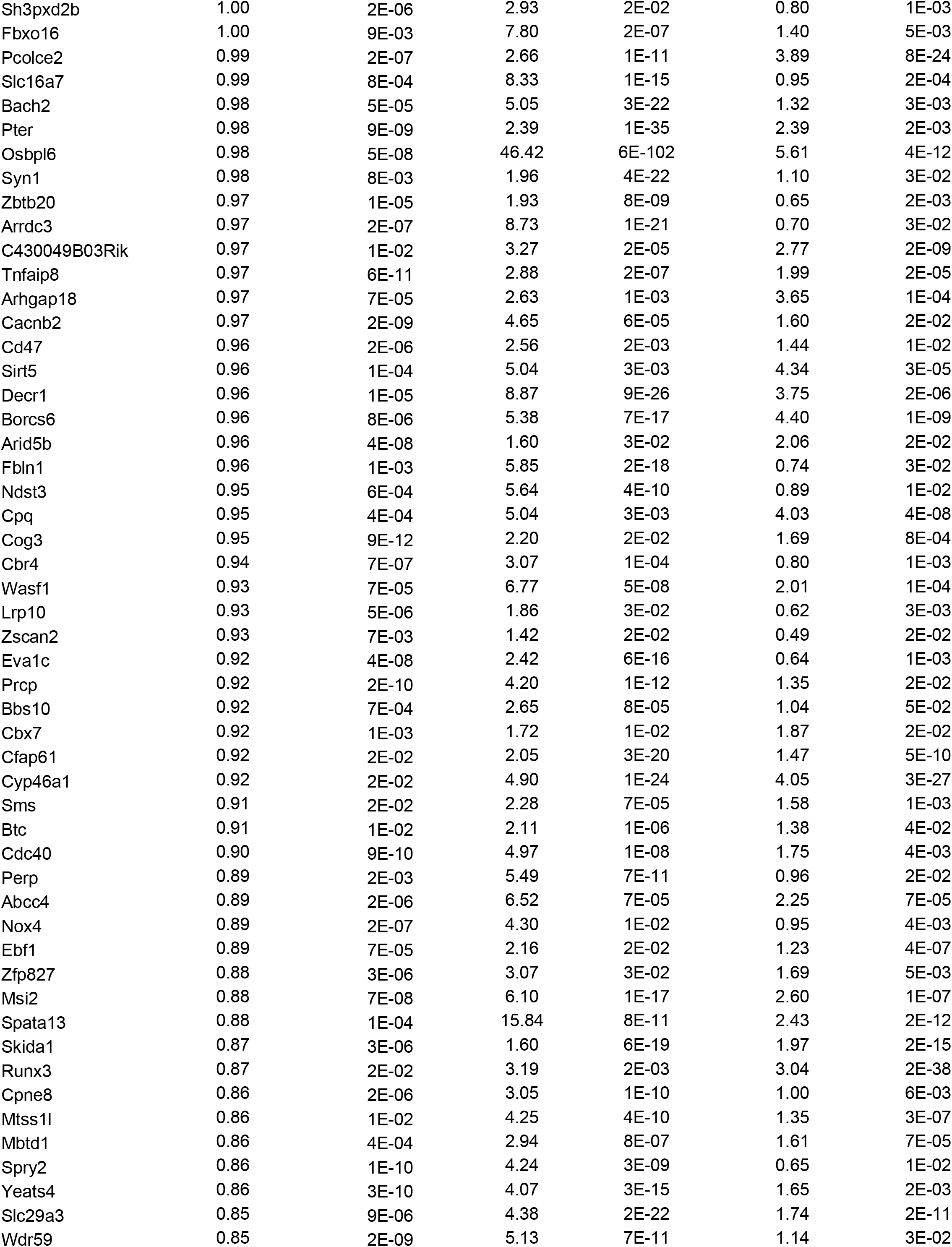
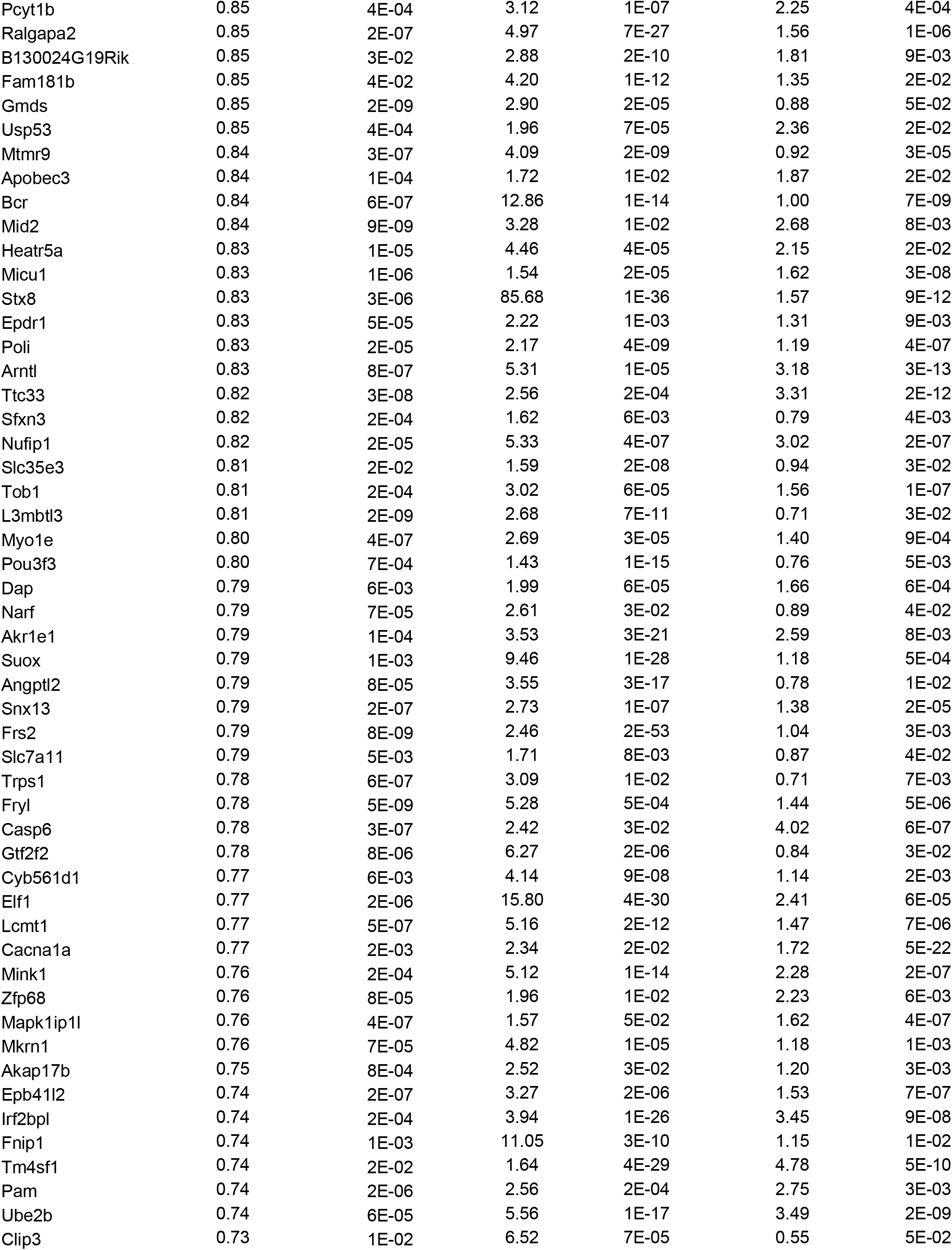
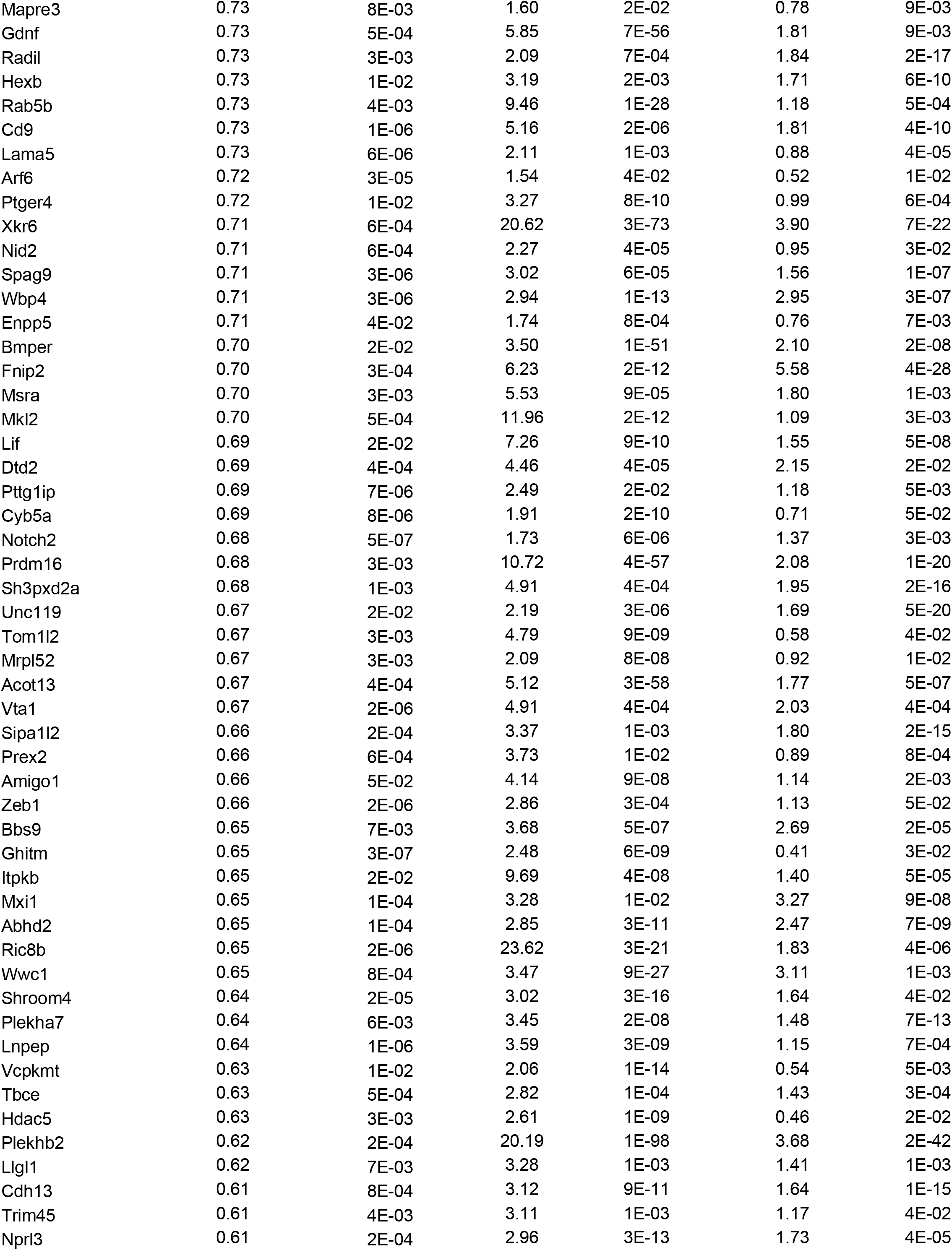
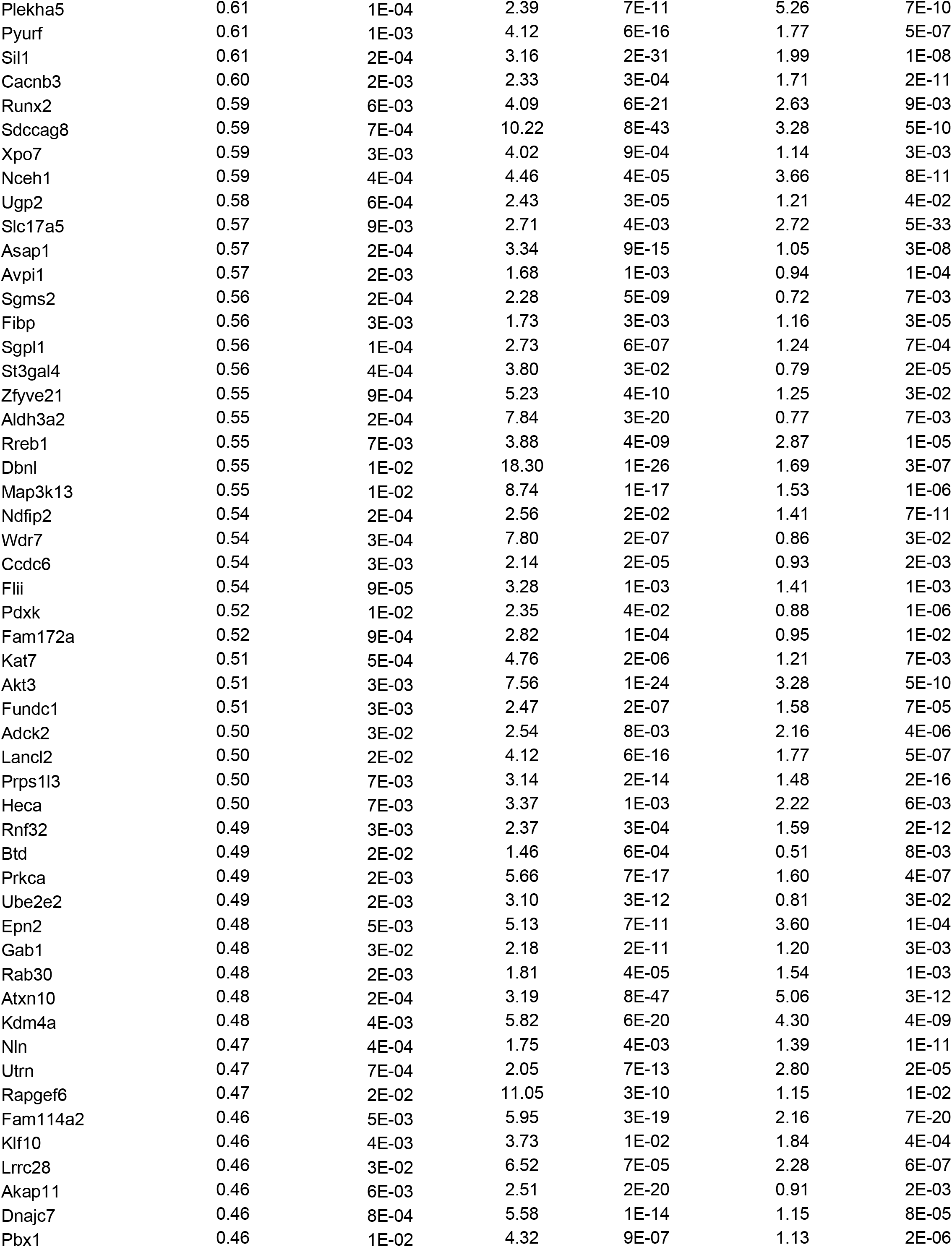
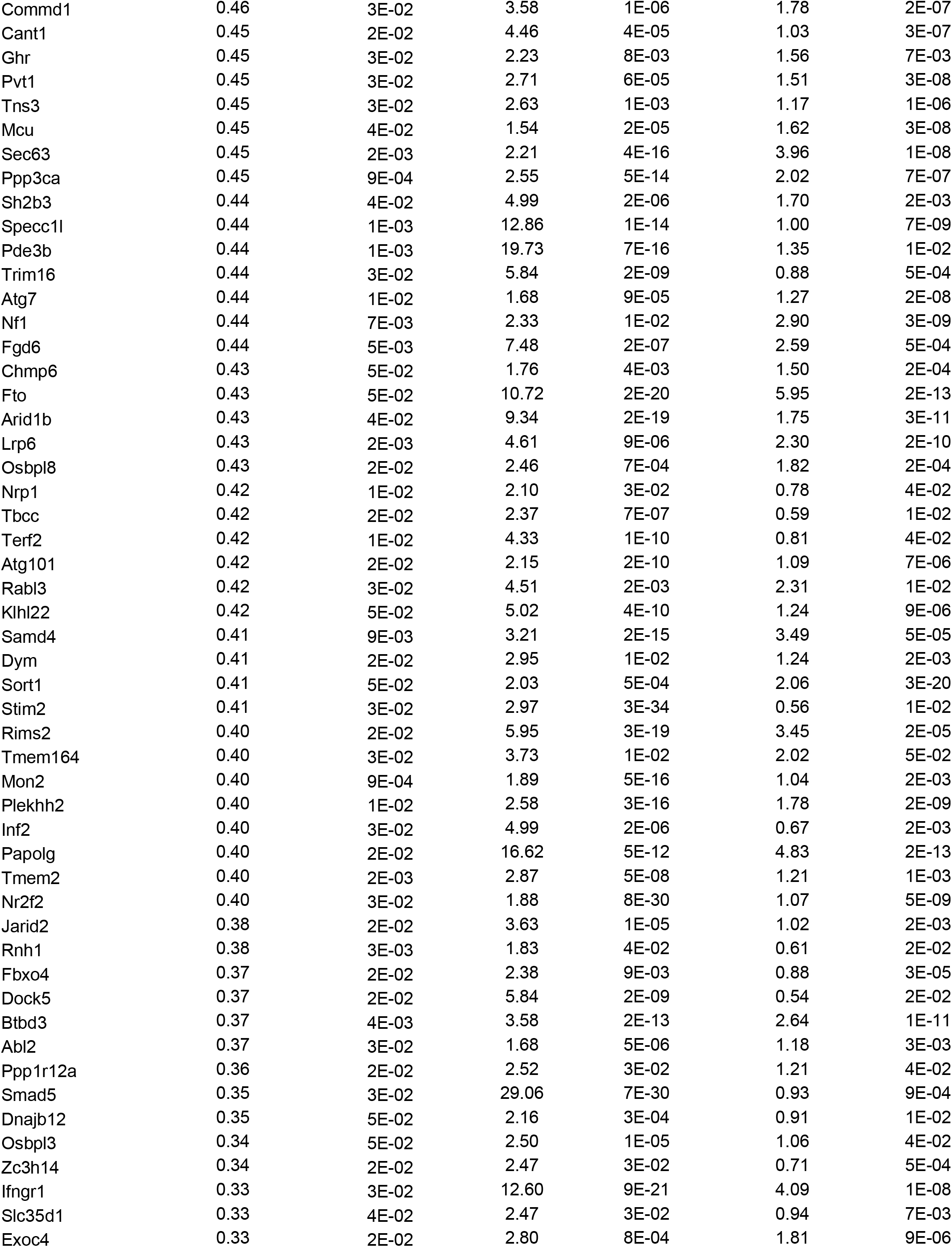
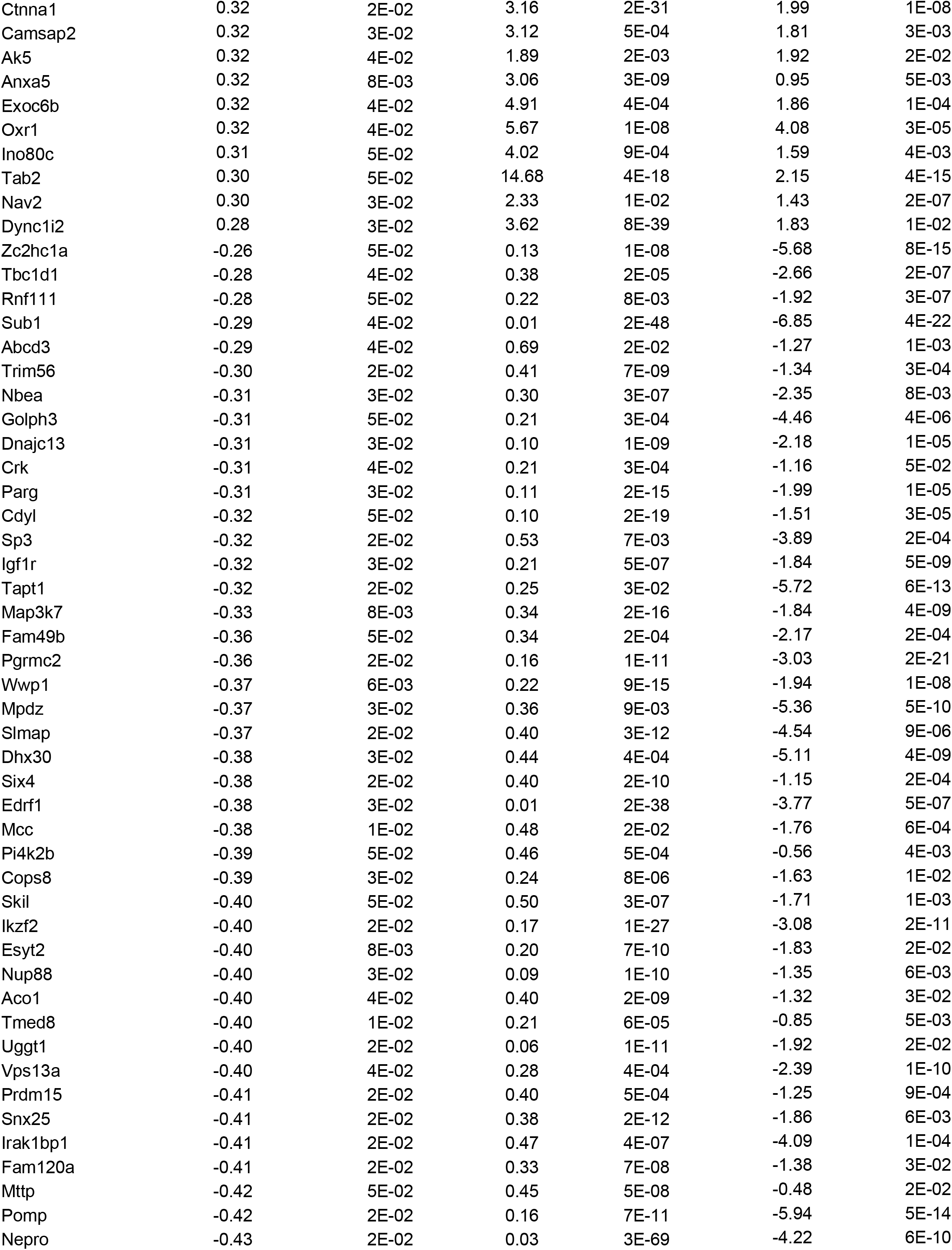
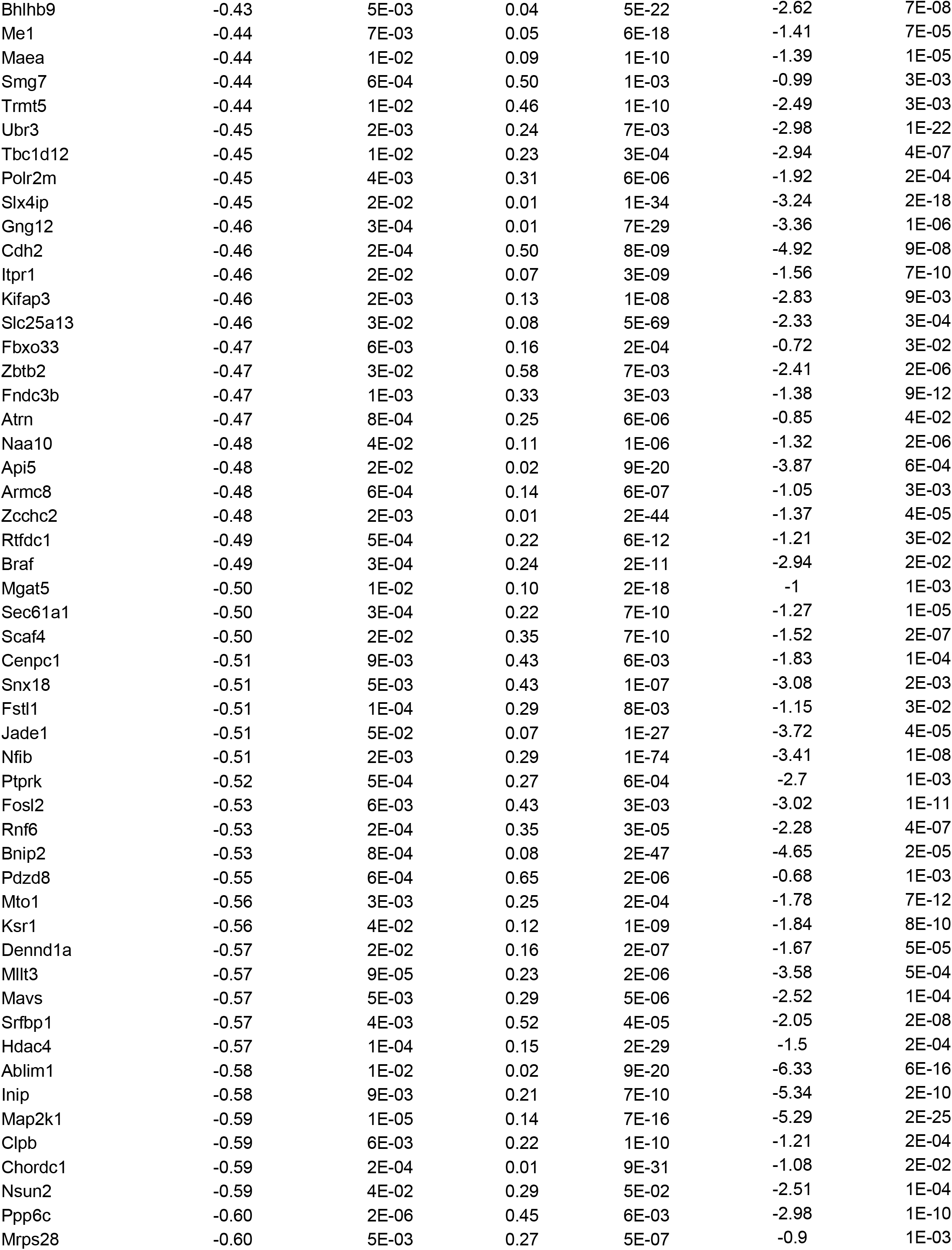
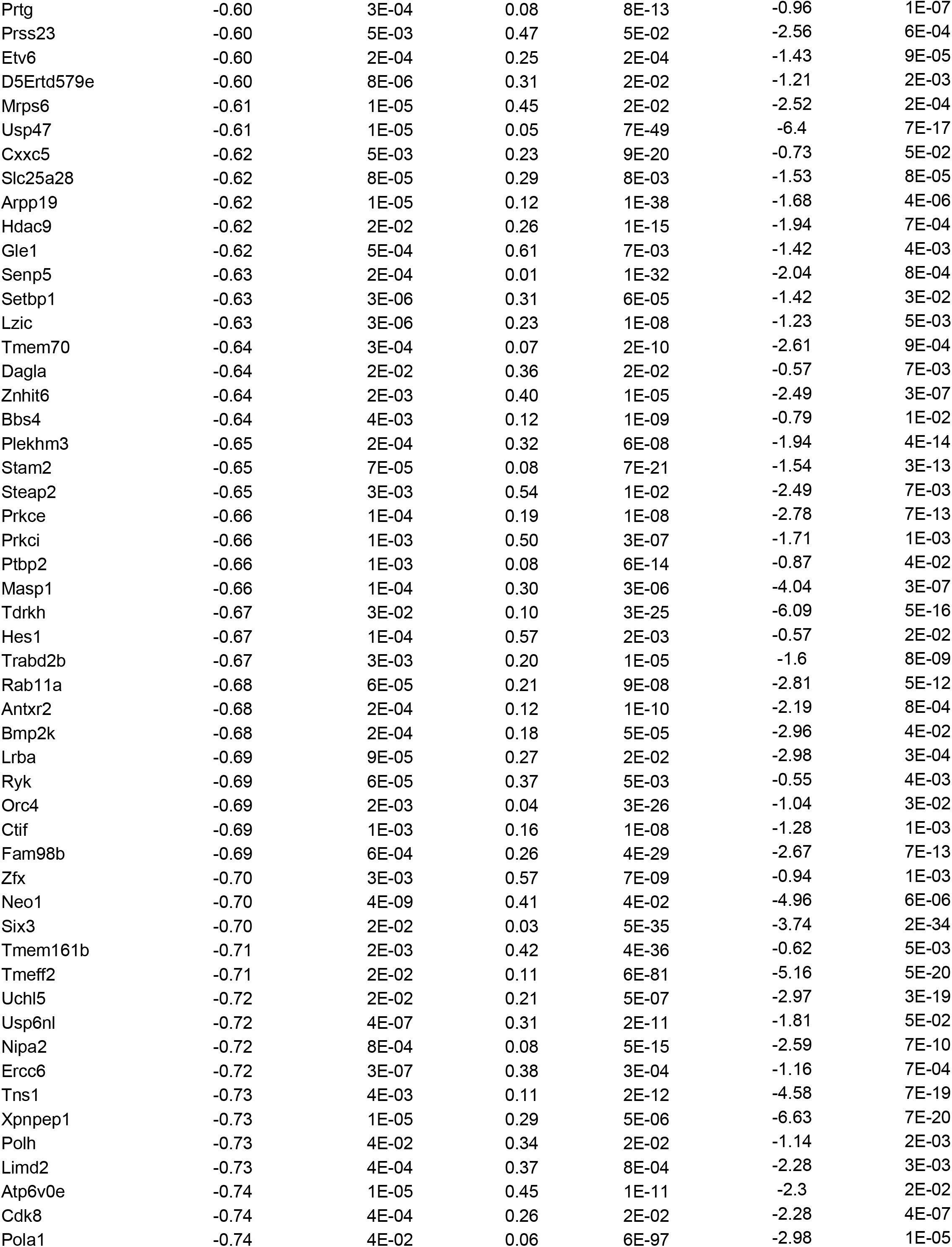
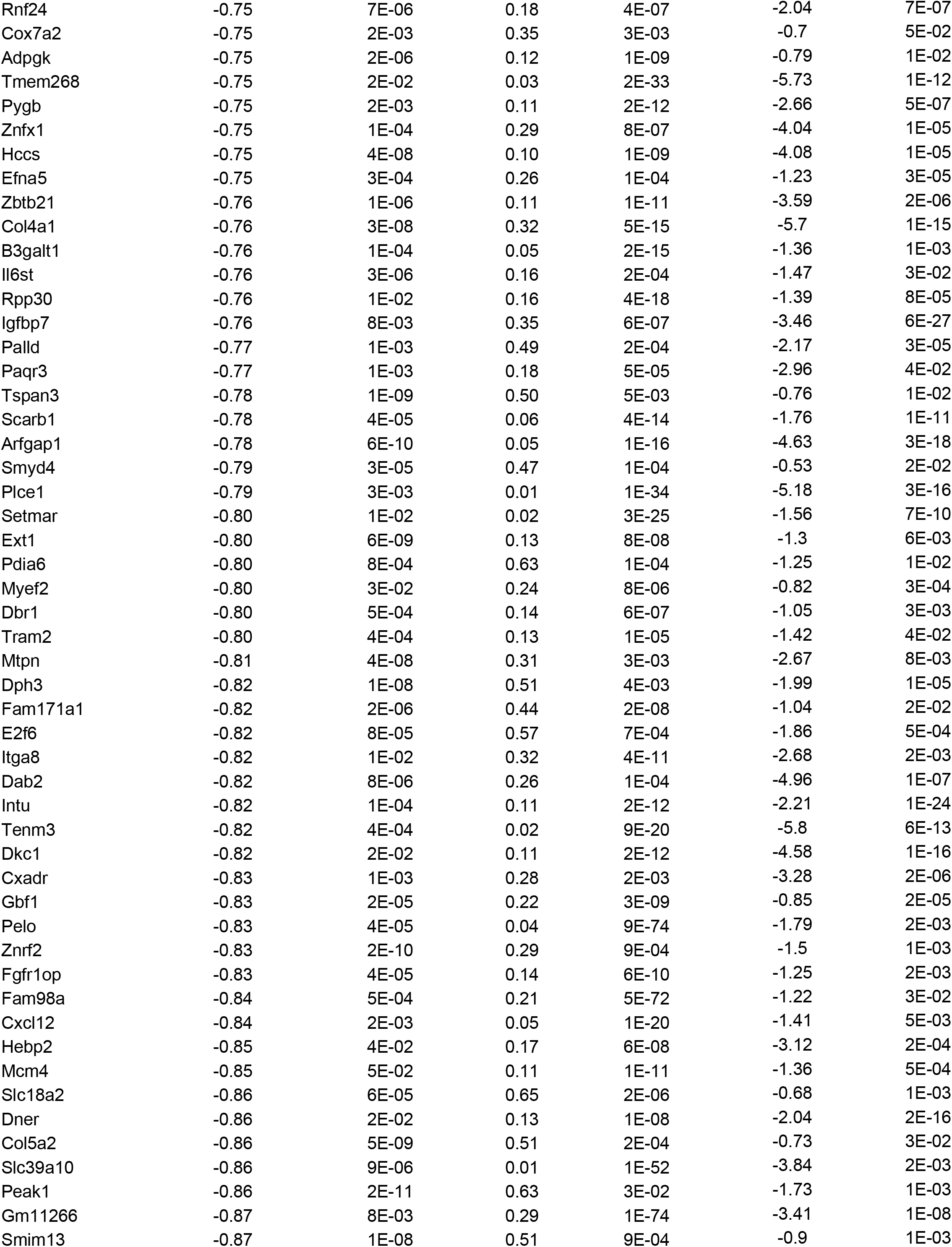
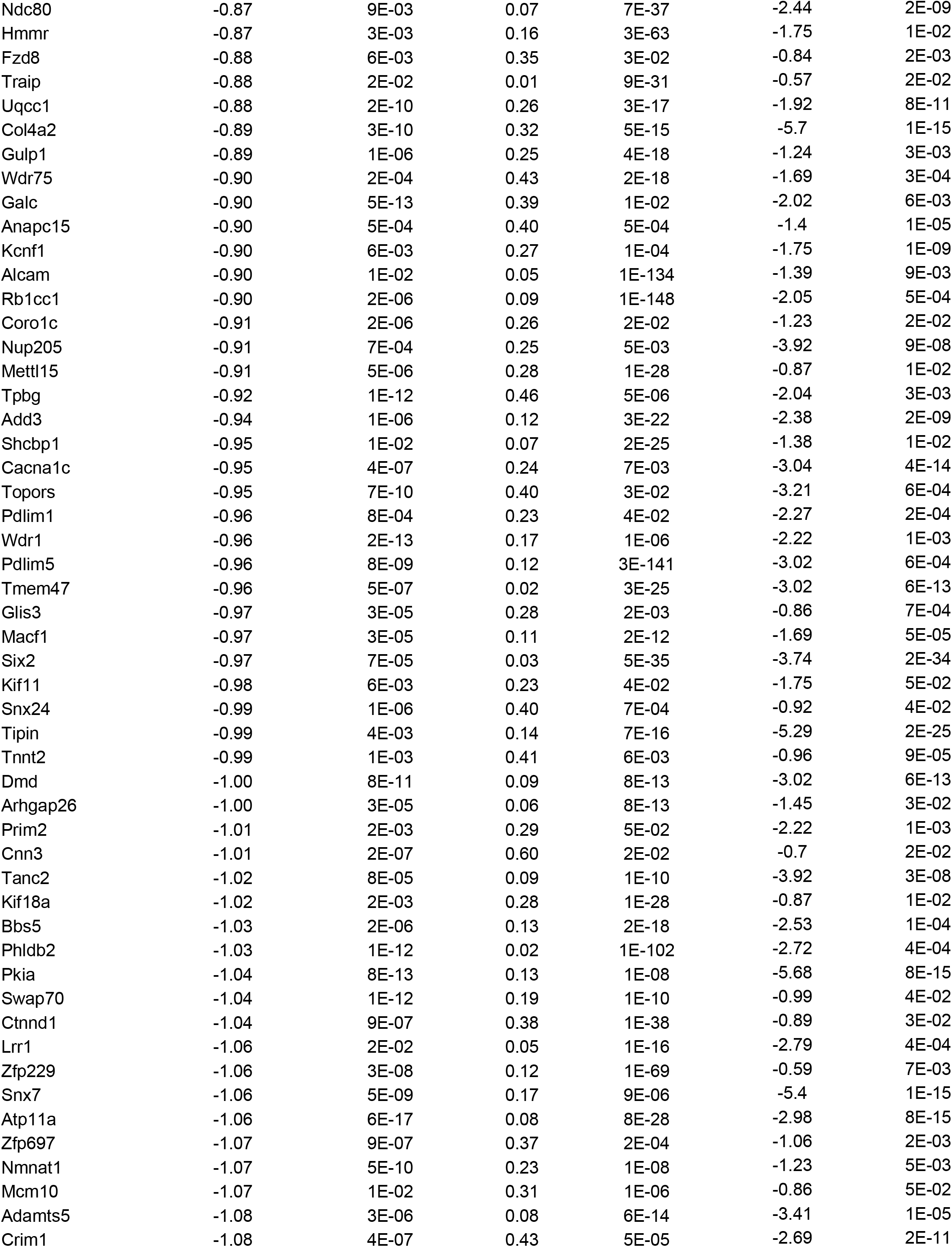
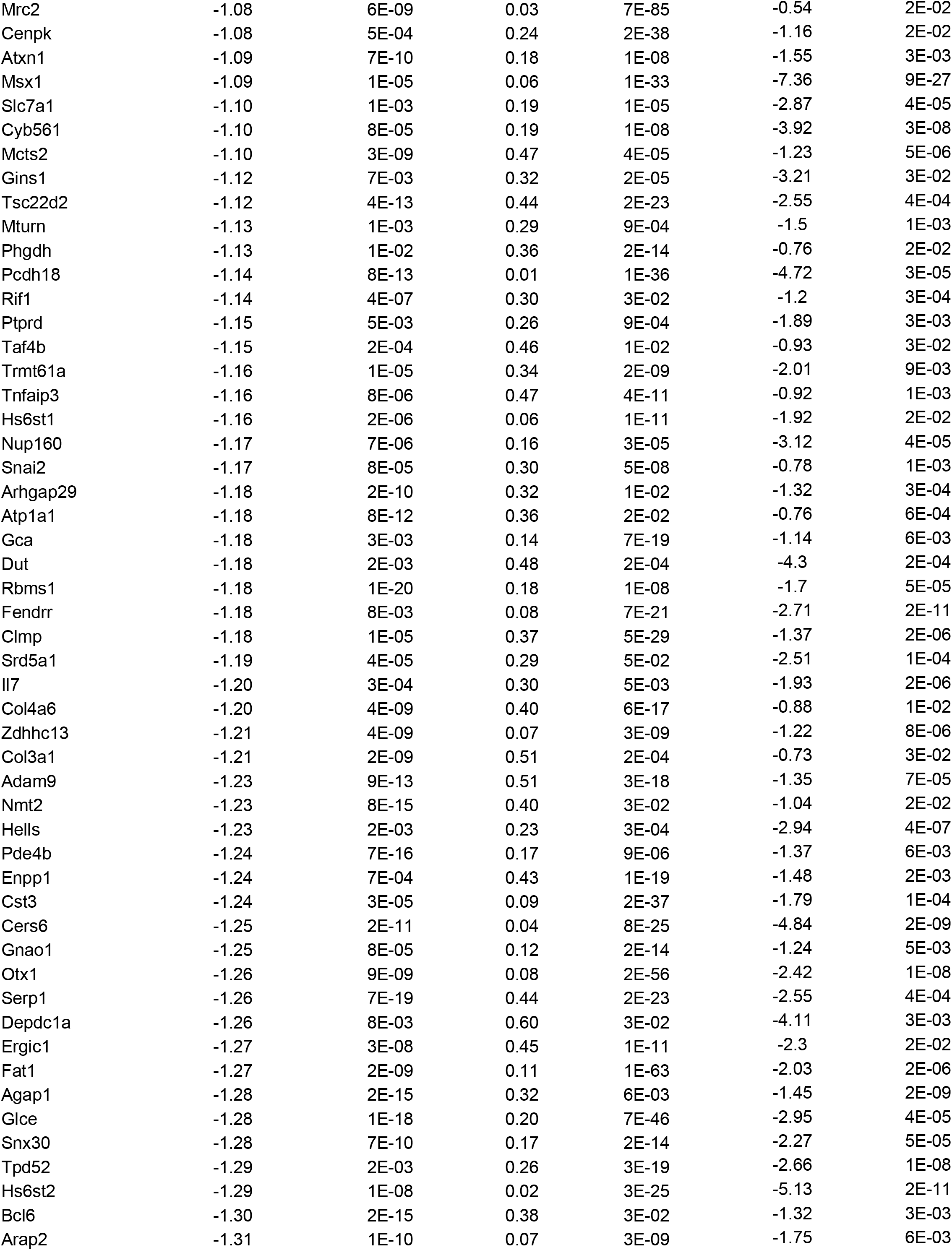
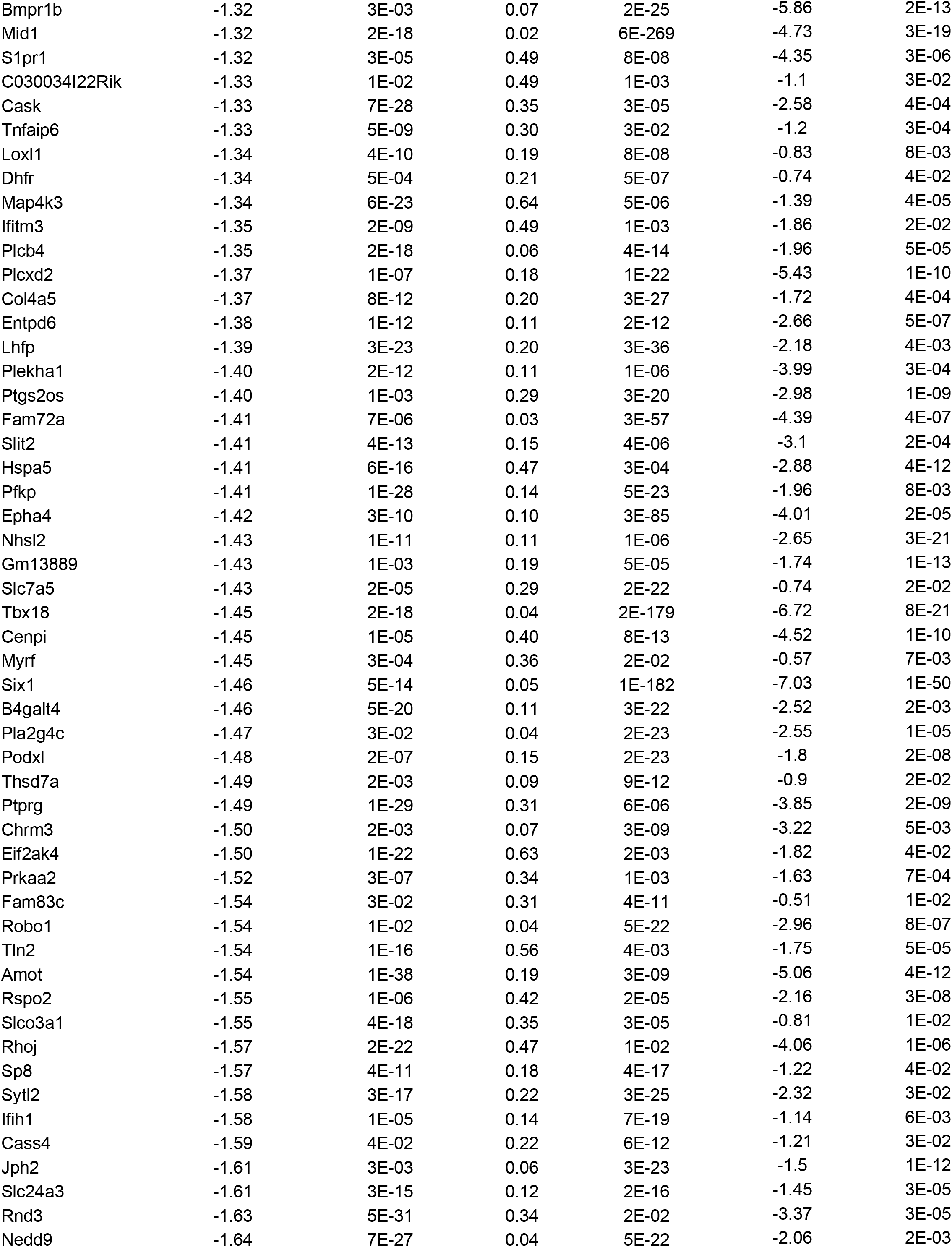
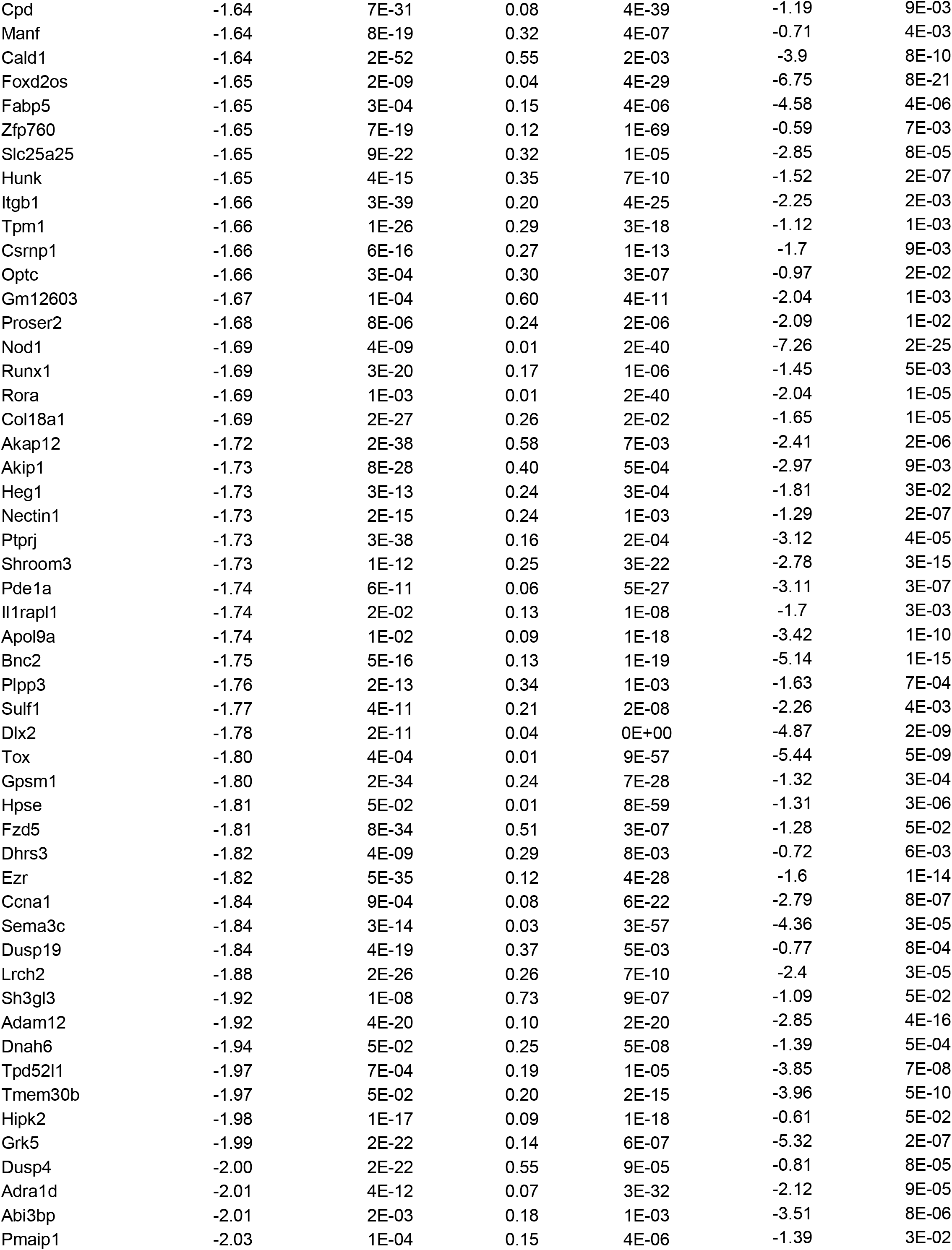
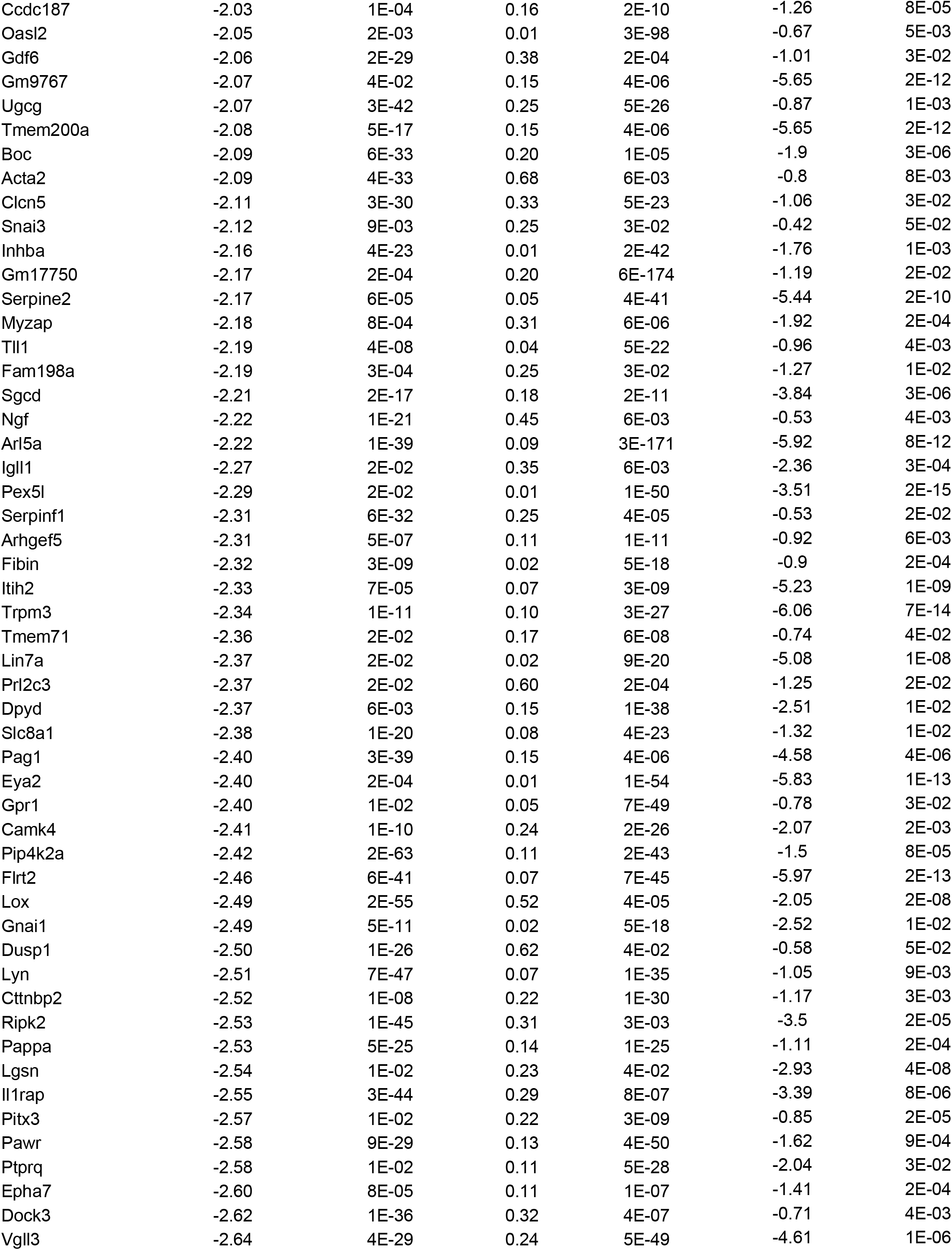
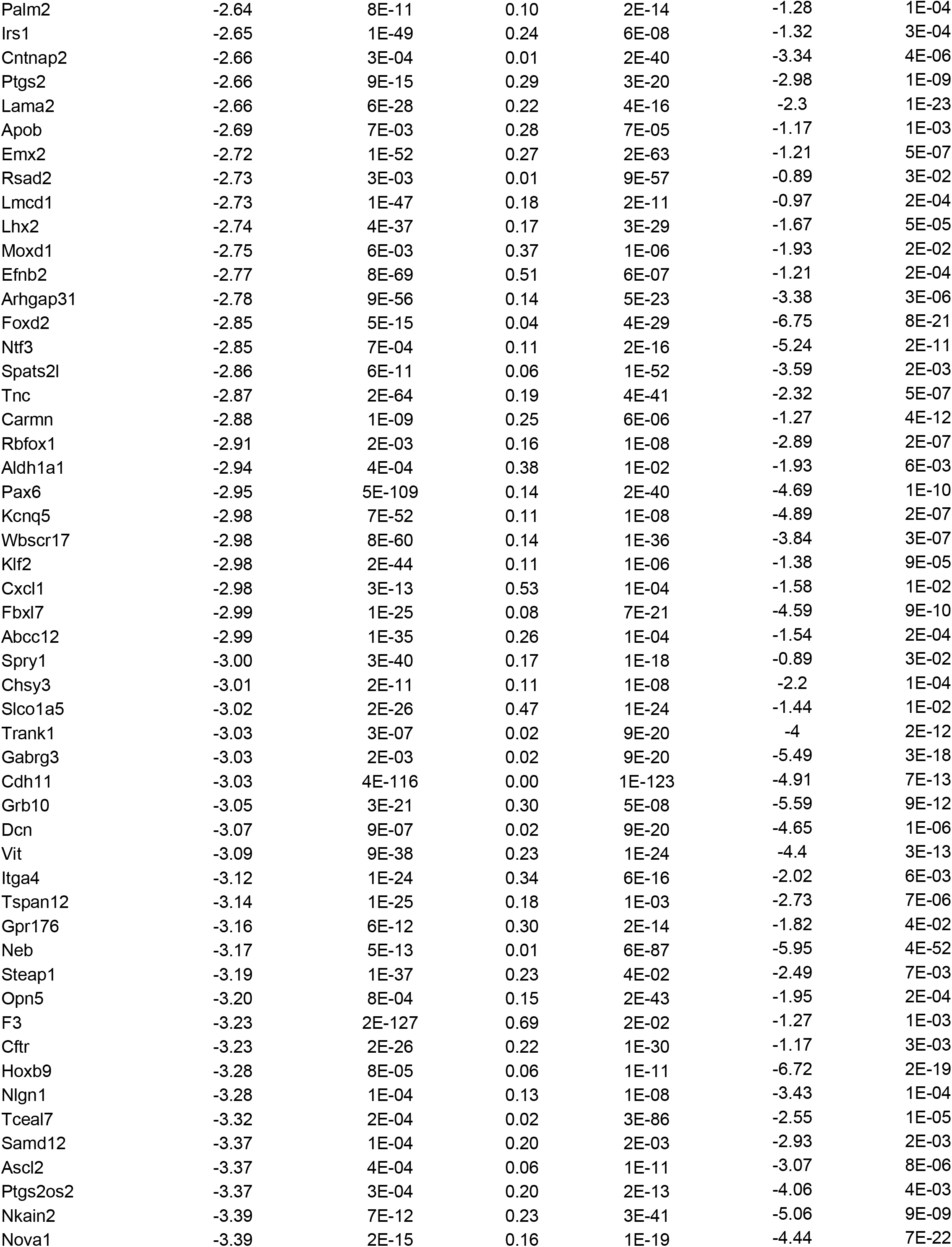
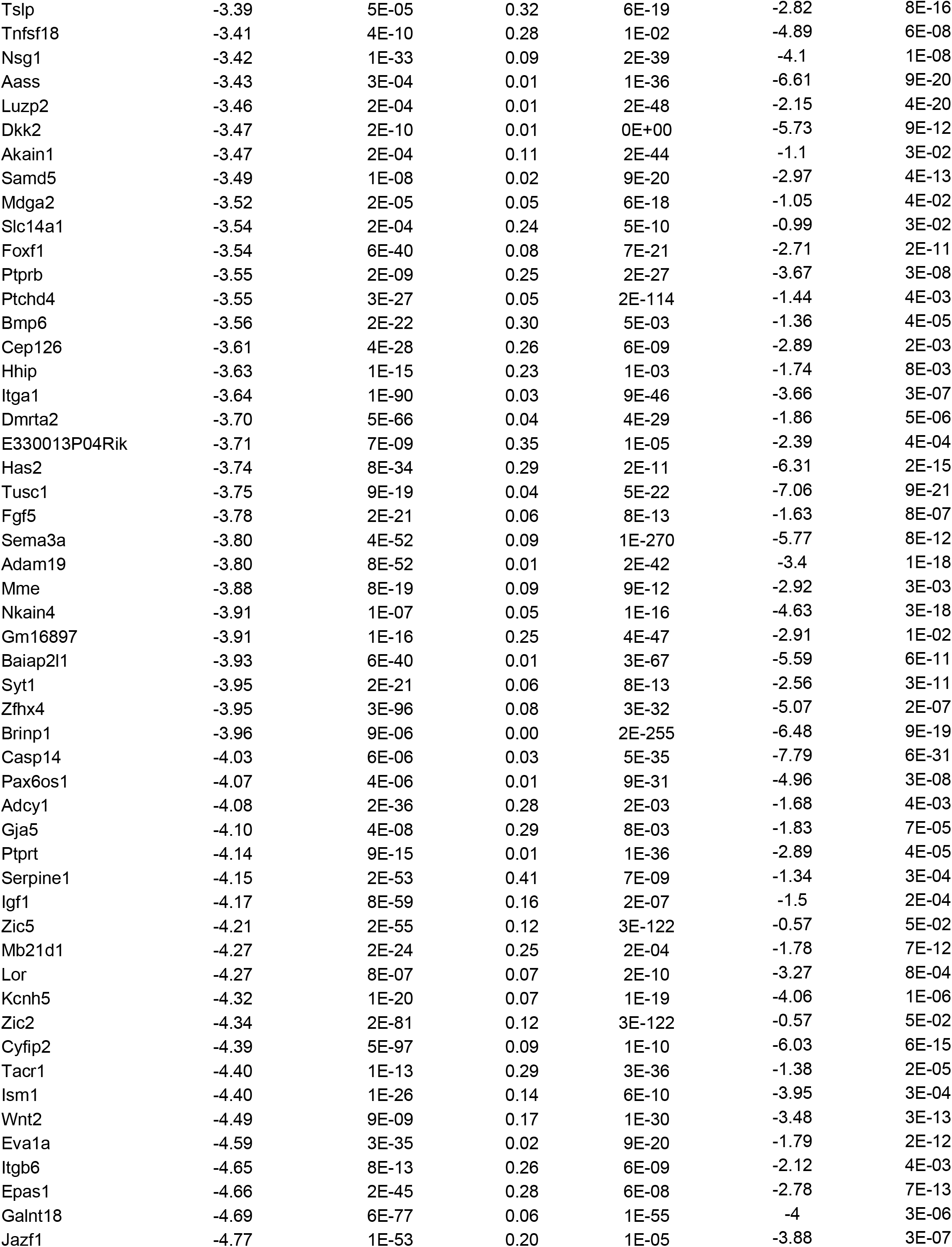
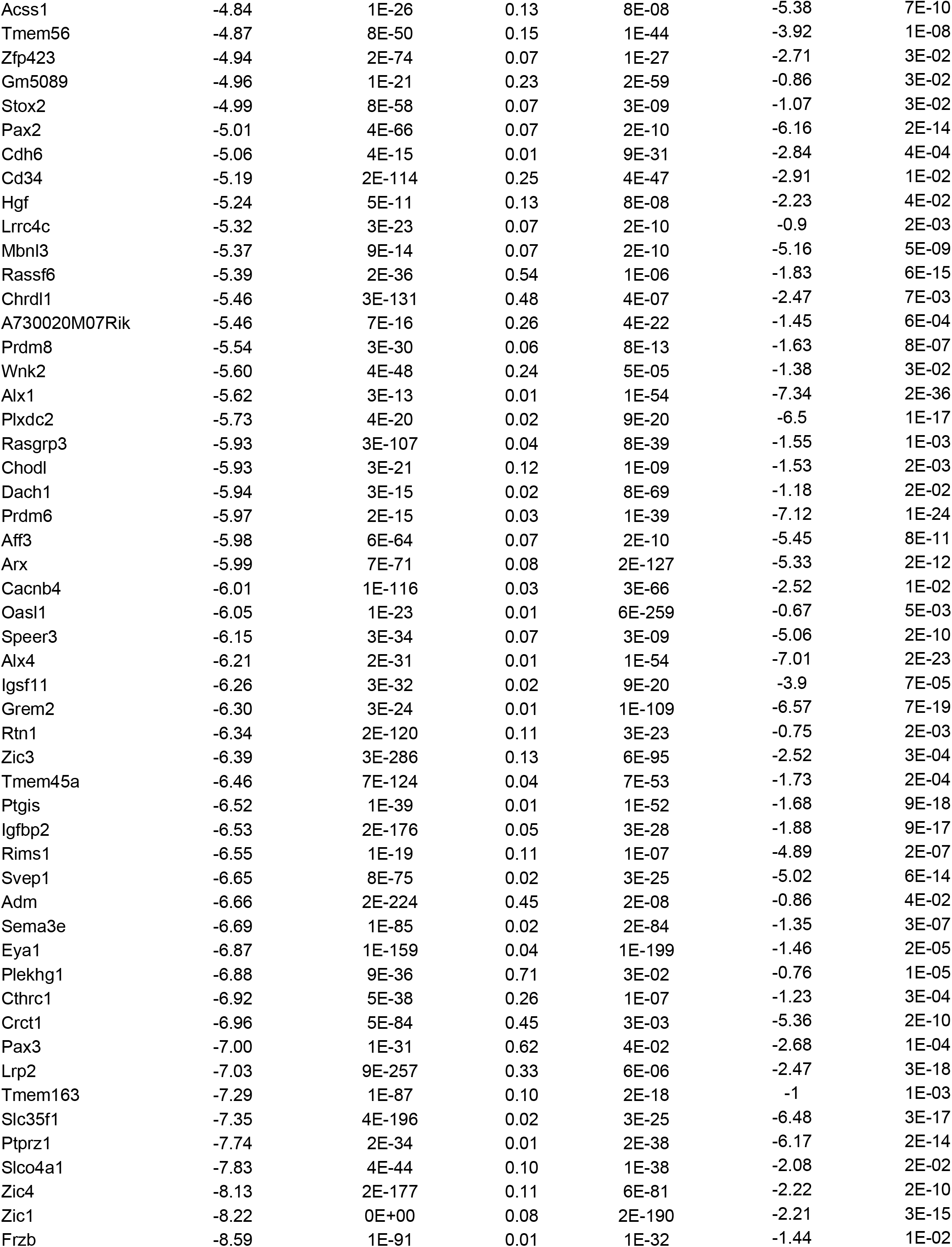
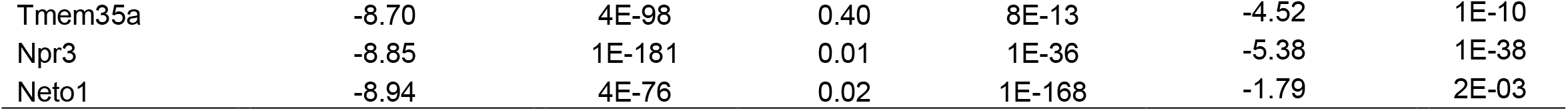
Differentially regulated sex-biased Brd4-bound genes in male and female GBM cells

**Table S3.** Genes with concordant gene expression in human GSCs and mouse GBM cells

| **Genes** | **Human GSCs**<br>**Linear Fold Change (M/F)** | **Human GSCs**<br>**Adjusted *p*-value** | **Mouse GBM Cells**<br>**Linear Fold Change (M/F)** | **Mouse GBM Cells**<br>**Adjusted *p*-value** |
| --- | --- | --- | --- | --- |
| DDX3Y | 1527.91 | 3.99E-271 | 6.37 | 6.92E-03 |
| APBB1IP | 59.00 | 4.01E-02 | 6.77 | 3.37E-20 |
| NSG2 | 22.47 | 1.40E-04 | 19.41 | 7.01E-14 |
| SYT13 | 18.46 | 6.74E-04 | 3.58 | 2.37E-04 |
| CRABP1 | 18.01 | 3.86E-04 | 11.18 | 8.59E-05 |
| GLT1D1 | 13.71 | 2.01E-03 | 13.94 | 1.11E-13 |
| GPR37L1 | 10.02 | 6.59E-03 | 5.50 | 1.54E-02 |
| RET | 9.98 | 3.28E-04 | 7.16 | 9.57E-05 |
| NRXN1 | 9.51 | 3.54E-02 | 4.70 | 9.20E-09 |
| DOC2B | 9.50 | 4.06E-03 | 5.42 | 4.08E-04 |
| CAMK1G | 7.97 | 4.31E-02 | 6.64 | 4.83E-03 |
| ALDH1L1 | 7.01 | 5.14E-03 | 14.61 | 2.63E-05 |
| GFRA1 | 6.97 | 2.40E-02 | 2.94 | 2.96E-06 |
| TEK | 6.68 | 4.10E-02 | 13.64 | 6.98E-09 |
| FUT9 | 6.52 | 3.28E-02 | 13.66 | 6.15E-10 |
| SCG3 | 6.00 | 2.34E-02 | 9.00 | 1.03E-04 |
| GPD1 | 5.83 | 4.57E-03 | 5.21 | 8.43E-17 |
| STXBP2 | 5.28 | 1.50E-02 | 3.82 | 4.74E-05 |
| NOTUM | 5.21 | 2.34E-02 | 5.37 | 1.33E-08 |
| TMEM151B | 4.66 | 1.50E-02 | 2.98 | 6.61E-03 |
| KCNAB1 | 4.58 | 3.09E-03 | 5.94 | 5.19E-20 |
| PHYHD1 | 4.51 | 4.10E-02 | 6.36 | 3.88E-05 |
| ATP8A1 | 4.09 | 6.83E-03 | 6.09 | 1.15E-30 |
| MYO7A | 4.09 | 9.18E-03 | 12.17 | 9.78E-20 |
| WNT10A | 4.03 | 2.05E-02 | 9.58 | 5.30E-05 |
| AOC3 | 3.33 | 1.70E-02 | 6.13 | 9.24E-10 |
| HID1 | 3.26 | 2.28E-02 | 8.08 | 4.06E-38 |
| PCDHB15 | 2.88 | 5.39E-03 | 3.69 | 4.72E-02 |
| ADHFE1 | 2.86 | 2.09E-02 | 10.56 | 1.01E-17 |
| RAPGEF5 | 2.85 | 4.76E-02 | 2.85 | 1.09E-05 |
| ITGA10 | 0.27 | 4.66E-02 | 0.25 | 1.86E-08 |
| CARMN | 0.20 | 3.19E-02 | 0.14 | 1.35E-09 |
| SERPINF1 | 0.10 | 1.70E-03 | 0.20 | 6.38E-32 |
| XIST | 0.0018 | 8.81E-112 | 0.0088 | 4.13E-49 |

**Table S4.**
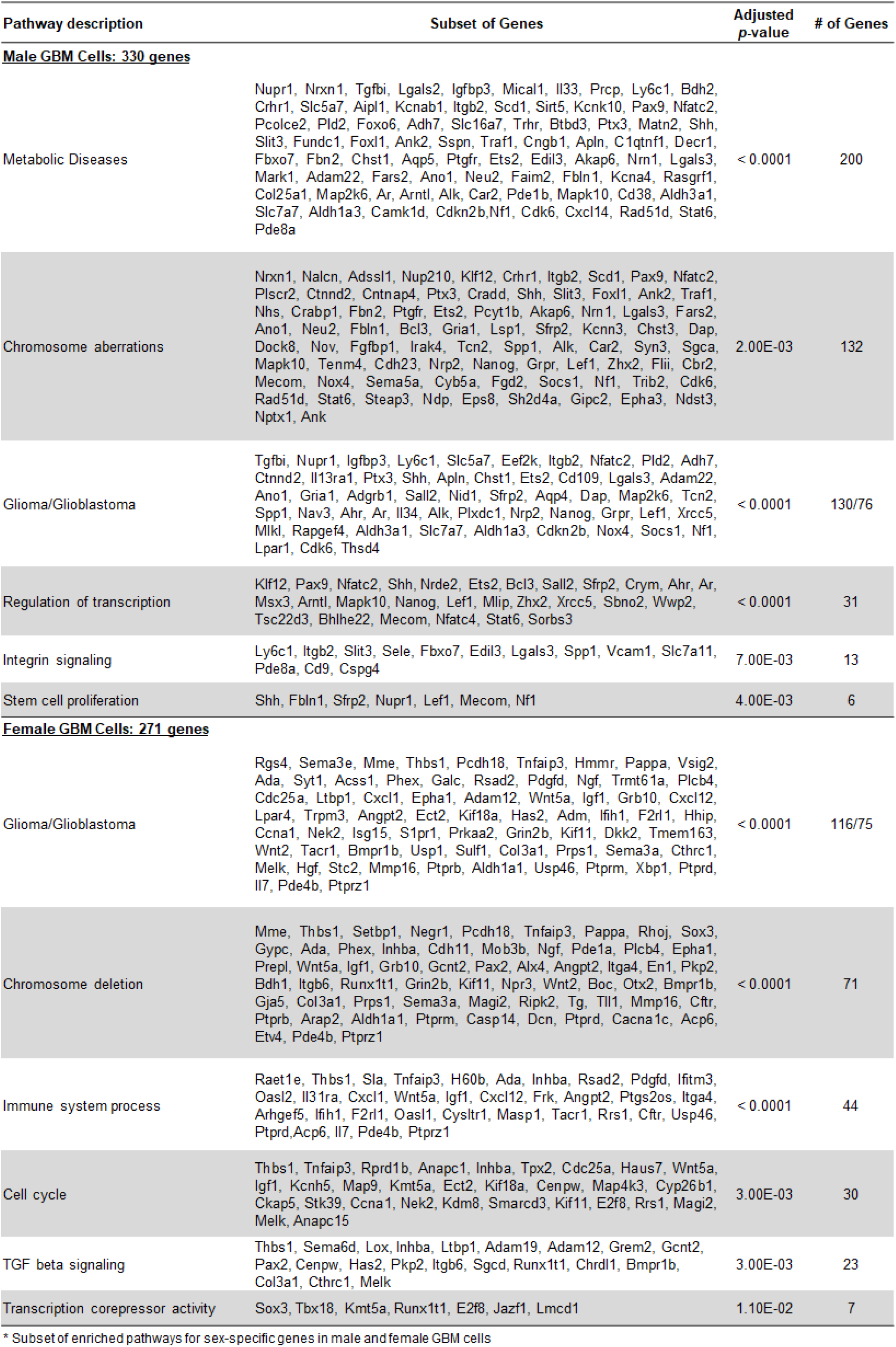
Pathway analysis for sex-biased Brd4-bound enhancer genes downregulated following JQ1 treatment in male and female GBM cells’

**Table S5.**
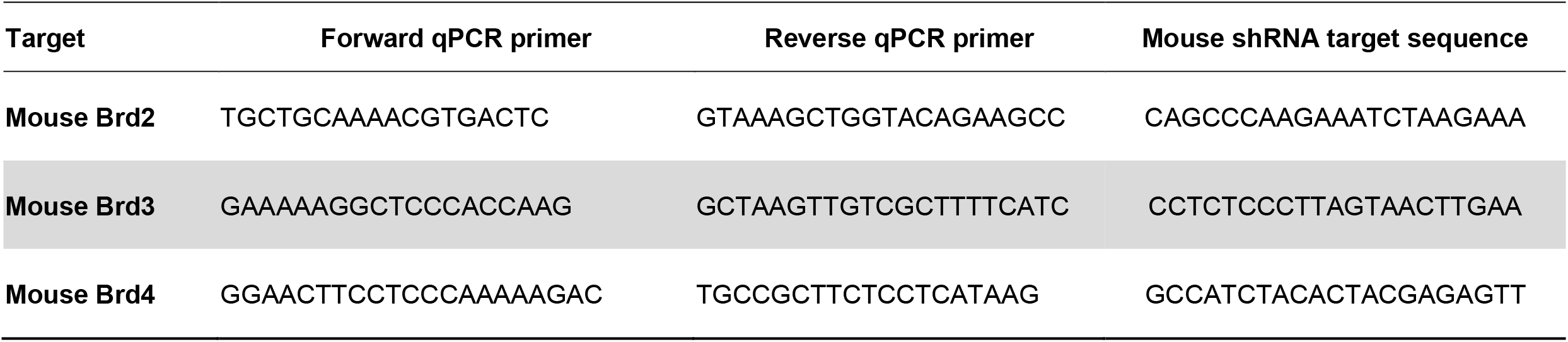
Real-time quantitative PCR primers and lentivirus shRNA target sequences

